# Potent Anti-Influenza Synergistic Activity of Theobromine and Arainosine

**DOI:** 10.1101/2024.10.13.618054

**Authors:** Hiya Lahiri, Eitan Israeli, Miriam Krugliak, Kingshuk Basu, Yelena Britan-Rosich, Tamar Ravins Yaish, Isaiah T. Arkin

## Abstract

Influenza represents one of the biggest health threats facing humanity. Seasonal epidemics can transition to global pandemics, with cross-species infection presenting a continuous challenge. Although vaccines and several anti-viral options are available, constant genetic drifts and shifts vitiate any of the aforementioned prevention and treatment options. Therefore, we describe an approach targeted at the virus’s channel to derive new anti-viral options. Specifically, Influenza A’s M2 protein is a well-characterized channel targeted for a long time by aminoadamantane blockers. However, widespread mutations in the protein render the drugs ineffective. Consequently, we started by screening a repurposed drug library against aminoadamantane-sensitive and resistant M2 channels using bacteria-based genetic assays. Subsequent *in cellulo* testing and structure-activity relationship studies yielded a combination of Theobromine and Arainosine, which exhibits stark anti-viral activity by inhibiting the virus’s channel. The drug duo was potent against H1N1 pandemic swine flu, H5N1 pandemic avian flu, aminoadamantane-resistant and sensitive strains alike, exhibiting activity that surpassed Oseltamivir, the leading anti-flu drug on the market. When this drug duo was tested in an animal model, it once more outperformed Oseltamivir, considerably reducing disease symptoms and viral RNA progeny. In conclusion, the outcome of this study represents a new potential treatment option for influenza alongside an approach that is sufficiently general and readily applicable to other viral targets.

## Introduction

Influenza presents a worldwide health challenge. Global estimates by the World Health Organization place the number of severe infections at 3–5 million, which leads to 290,000– 650,000 deaths annually (1, 2). Therefore, it is no surprise that before the COVID-19 pandemic, it was the leading cause of mortality in the Western world from infectious diseases (3). The etiological agent of influenza is the eponymous Influenza virus, identified by Shoppe in 1931 as the cause of swine flu (4–7) and by Smith, Andrewes, and Laidlaw in 1933 as the agent behind the human disease (8). As a member of the *Orthomyxoviridae*, the virus is a negative-sense, single-stranded, segmented RNA virus. The family contains several genera, of which only the *Alphainfluenzavirus* and *Betainfluenzavirus* have members that pose a health risk. Moreover, only the Influenza A virus has been known to cause pandemics and, therefore, is the focal point of our study.

There are several prevention and treatment options for influenza: Annual vaccinations have been available for some time, but their efficacy is limited between 40% and 60% (1) due to the low fidelity of its RNA polymerase that causes a constant drift and shift in the virus’s genome. The principal antigens are the hemagglutinin and neuraminidase glycoproteins, of which there are 16 and 9 serotypes, respectively (not including bat Influenza A-like viruses(9)). Consequently, their combination is used as a primary descriptor of the strain, and in the current study, we focus on the two most significant health threats: H1N1 swine flu and H5N1 avian flu.

Anti-viral agents are also available against the Influenza virus, targeting several of the pathogen’s proteins (for a recent review, see (10)). The most common agents in current use are neuraminidase inhibitors such as Oseltamivir, Zanamivir, Peramivir, and Laninamivir. More recently, inhibitors were approved against the viral RNA polymerase and its complex (Favipiravir and Baloxavir) and the pathogen’s fusion machinery (Umifenovir). However, the first anti-influenza agents were the aminoadamantanes, identified in 1964 (11). Mutations in M2 conferred resistance to said drugs, thereby singling the protein as the target within the virus (12). In 1992, Lamb and colleagues, in a landmark study, showed that M2 is a H^+^ channel that is blocked by aminoadamantanes, thereby providing a molecular mechanism of their activity (13). Finally, resistance has developed against all aforementioned drugs (10), and is particularly prevalent against aminoadamantanes, rendering their utility obsolete (14).

The discovery of an ion channel coded by the Influenza virus led to the identification of many such proteins, collectively termed viroporins (15). They are often characterized by their small size and oligomeric nature, with varying impacts on the viral infectivity cycle. In Influenza, M2 was shown to be essential to the pathogen since it allowed the concomitant acidification of the viral lumen upon endocytosis and release of the viral RNP from the matrix (13).

Considering the fact that channels, as a family, are excellent drug targets, and there is a dearth of curative anti-influenza agents, we sought to identify novel blockers against the M2 protein. In particular, we searched for blockers that will be active against aminoadamantanes and aminoadamantane-resistant channels. To that end, we employed a combined approach of bacteria-based assays, *in cellulo* assays, and structure-activity analyses that led to successful animal experimentation. Finally, this approach yielded a potent anti-influenza agent and represents a general route applicable to many other viruses.

## Materials and Methods

### Bacteria-based channel assays

Three individual bacteria-based assays were used to examine the channel activity of the viroporin and blocker activity thereupon. The MBP (maltose binding protein) fusion purification system (New England BioLabs; Ipswich, MA, USA) was used, whereby Influenza A (Singapore/1/1957(H2N2)) M2 was expressed as a chimera by fusing it to the carboxy-terminus of the maltose binding protein to ensure proper membrane reconstitution (16). The aminoadamantane-resistance mutation, S31N (17), was obtained by site-directed mutagenesis of the above Singapore strain, which is aminoadamantane-sensitive (18). Sequences of these and additional M2 proteins are given in Supporting Fig. 1.

**Fig 1.**
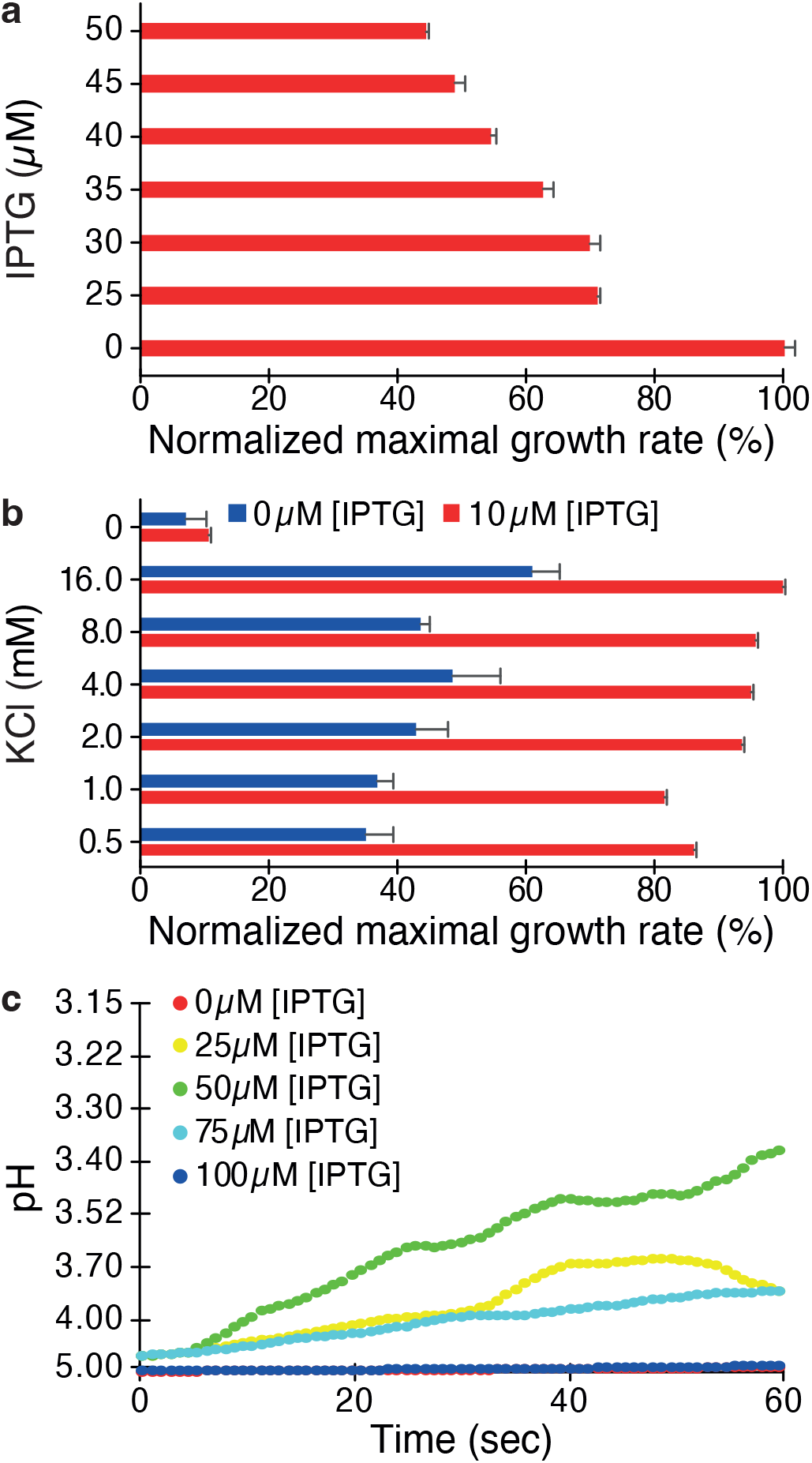
Three bacteria-based assays to assess the channel activity of the Influenza M2 protein: (a) Negative assay, (b) Positive assay, and (c) pH-dependent fluorescence (pHlux) assay. Note that in the negative assay, protein expression retards bacterial growth; in the positive assay, protein expression increases growth; and finally, in the fluorescence-based essay, protein expression changes bacterial pH.

#### Negative assay

DH10B cells (Invitrogen; Carlsbad, CA, USA) containing Influenza-M2 and S31N-M2 plasmids were grown overnight in lysogeny broth (LB) medium. Following a 50-fold dilution, the culture was set to grow until its OD600 reached a value of 0.2. Subsequently, varying concentrations of Isopropyl-*β*-D-1-thiogalactopyranoside (IPTG) ranging from 25 μM to 50 μM at 5 μM intervals were employed to evaluate the impact of protein expression on bacterial growth. Bacteria grown in 96-well plates were incubated in a multiplate reader (LogPhase 600 from BioTek; Santa Clara, CA, USA) at 37°C for 16 hrs, and readings were taken at 15 min intervals.

#### Positive assay

K^+^-uptake-deficient bacteria (LB650: *ΔtrkG, ΔtrkH*, and *ΔkdpABC5*) were used in this assay. These particular bacteria are incapable of growth in regular LB medium, unless they express a channel capable of potassium transport, or if the growth medium contains a high concentration of potassium (19). Overnight growth of bacteria was conducted in LBK medium, which was identical to LB expect that KCl replaced NaCl. Different concentrations of KCl and Isopropyl-*β*-D-1-thiogalactopyranoside were used to gauge the impact of the channel’s activity on the bacterial phenotype.

#### pHlux assay

The pHlux assay (20) is based on LR1 bacteria that express a chromosomal copy of a pH-sensitive GFP (21). The bacteria were grown overnight in the same conditions as DH10B cells. Subsequently, the culture it is diluted to 1:50 and again grown until the OD600 reaches a values of 0.2. Different concentrations of Isopropyl-*β*-D-1-thiogalactopyranoside are used to induce protein expression for one hour and cells are diluted to an OD600 of 0.3 and pelleted down at 3500 g for 10 min. The cells are then washed twice and resuspended in McIlvaine Buffer(22) which contains 200 mM Na2HPO4 and 0.9% NaCl adjusted to pH 7.6 with 0.1 M citric acid. The spectroscopy is undertaken in 96-well plates (Nunclon f96 Microwell Black Polystyrene, Thermo Fisher Scientific; Waltham, MA, USA), and in each well 200 μL of cell suspension was added along with 30 μL of buffer. 70 μL of citric acid (300 mM, 0.9% NaCl) are then added to each well by a liquid handling system (Tecan; Männedorf, Switzerland) and readings were carried out at two excitations of 390 nm and 466 nm with emission held at 520 nm (21). The proton flow was calculated from the ratio of two excitations as described previously (20).

### Channel blocker screening

A library of 2,839 compounds was purchased from MedChem Express (HY-L035, Monmouth Junction, NJ, USA) and used for screening against the aminoadamantane-sensitive and S31N (aminoadamantane-resistant) Influenza virus M2 channels. The screening employed the negative assay with a 100 μM drug concentration with a total amount of DMSO of 2%. Protein expression was achieved by adding 45 μM of Isopropyl-*β*-D-1-thiogalactopyranoside which retarded growth by approximately 50% (Fig. 1). Bacteria that received only DMSO were designated as negative control. Two metrics were recorded: growth rate and final growth density. Drugs that were able to enhance bacterial growth by a certain threshold were subsequently checked in duplicate.

Successful compounds were also subjected to the positive genetic assay where they were examined at 10 μM Isopropyl-*β*-D-1-thiogalactopyranoside and 8 mM KCL conditions. Finally, compounds that passed both the negative and positive assays were subjected to the pHlux assay.

### *In-vitro* studies

#### Cell culture

Madin-Darby canine kidney (ATCC MDCK NBL-2) cells were maintained in Dulbecco’s Modified Eagle’s Medium (DMEM) (Biological Industries; Beit Haemek, Israel), supplemented with 10% fetal bovine serum, 2 mM L-Glutamine, 10 IU/mL Penicillin, 10 μg/mL streptomycin, and Biomycin-3 (Biological Industries).

#### Virus culture and infection

Influenza A Virus A/Wisconsin/629D02452/2009 (H1N1)pdm09 was obtained through BEI Resources, NIAID, NIH (NR-19810). Influenza A Virus A/Puerto Rico/8/34 was acquired from ATCC (VR-1469). H5N1, A/Israel/975/2023 was collected from a peregrine falcon and obtained from the Israeli veterinary service. The M2 sequences from the three viruses are given in Supporting Fig. 1. Identical aliquots of virus stock were prepared from the mother stock. For infection, 1:1000 diluted sub-stocks were prepared from each aliquot. Subsequent infection of MDCK cells was carried out in DMEM containing 0.3% Bovine Serum Albumin, (Sigma Aldrich) and 3 μg/ml TPCK treated trypsin (Sigma Aldrich) and further incubated for 48 hrs at 37°C in a 5% CO2 atmosphere. All infection experiments were performed in a BSL-3 facility.

#### Cell viability and cytotoxicity assays

MDCK cells were cultured in a 96-well plate with 200 μL of medium at density of 15,000 cells per well and grown overnight. The dilutions of tested compounds were prepared in DMEM with 0.3% BSA, 3 μg/ml TPCK-treated trypsin, and 50 μL of said solution was added to the cells. The effects of the drugs on the viability of the cells were assessed at 48 hrs post-treatment using CellTiter 96 Aqueous Non-Radioactive Cell Proliferation reagent (Promega; Madison, WI, USA). To examine the effect of different drugs, cells were infected with the Influenza virus at a multiplicity of infection (MOI) of 0.3 for two hours followed by treatment with various drugs. Infection with the PR8 strain employed 3 ug/ml of TPCK treated trypsin and an MOI of 1. Each compound concentration was tested in triplicate and each assay plate contained the following controls: no cells (background control), cells treated with medium (mock infection for normalization), infected/untreated cells, and infected/solvent-treated cells (infection control).

At two days post-infection, anti-viral drug efficacies were assessed by the CellTiter 96 Aqueous Non-Radioactive Cell Proliferation reagent (Promega) for 3 hrs at 37°C in a 5% CO2 atmosphere. Reactions were stopped and the virus was inactivated by adding 30 μL of 4% formaldehyde. Absorbance was measured at 492 nm using a Tecan plate reader Finally, the data were normalized to the mock-infected control, after which inhibition metrics values were calculated by fitting the data to a Monod equation (half-saturation coefficient or *Ks*).

#### Viral load quantification

To check viral load from each experiment, the supernatant was collected after 48 hrs of infection and RT-qPCR was performed. Viral RNA was extracted from the supernatant using the Aurum™ total RNA miniKit (Bio-Rad; Hercules, CA, USA). Subsequently, cDNA was synthesized from the extracted viral RNA according to the qScript cDNA synthesis kit protocol (Quanta bio; Beverly, MA, USA).

Detection of Influenza virus RNA was done with primers specific to the M gene: forward primer 5’-CGCTCAGACATGAGAACAGAATGG-3’ and reverse primer 5’-TAACTAGCCTGACTAGCAACCTC-3’ (Integrated DNA Technologies; Coralville, IA, USA). The reaction was conducted in a 96-well plate, with 2 μL of cDNA, 1× iTaq Universal SYBR green supermix (Bio-Rad) and 200 nM forward and reverse detection primers to a total volume of 20 μL/well.

For a standard curve from purified RNA, a stock concentration of 10^6^ copies/ml was used and cDNA was prepared and then serially diluted with elution buffer. Thermal cycling was performed using StepOnePlus™ Real-Time PCR System with StepOne™ Software Version 2.3 (Carlsbad, CA, USA).

#### Structural similarity studies

Chemicals that are structurally similar to the hit compounds (obtained from the initial channel screening and *in-vitro* studies) were searched to examine their anti-viral activity. In this regard, Tanimoto similarity searches with a value of 85% and above were performed with each hit compound to get several similar chemicals available commercially.

#### *In-vivo* studies

All procedures involving animals took place at the Authority for Biological and Biomedical Models of the Hebrew University of Jerusalem, accredited by the Israeli Council for Experiments of Animal Subjects and by the Association for Assessment and Accreditation of Laboratory Animal Care, International (AAALAC). The anti-viral efficacy experiments were conducted at the national biosafety Level 3 biocontainment unit. Finally, all experiments were conducted under IACUC-approved protocols (MD-23-17229-5 and MD-23-17159-5).

All animal experiments employed six-week-old male BALB/c mice. Throughout the experiment, a scoring table was used to monitor the condition of the animals alongside recording body weights daily. Death is not an endpoint in the experiment. Rather, in case of more than 20% of body weight loss, euthanasia was performed. In addition, a summation of the clinical signs according to the General Mouse Scoring Table (see Supporting Tables S1 and S2), was recorded at least twice daily. The animals were monitored thrice a day if the score is higher than 12. An early withdrawal point and euthanasia occured when a total score of 15-16 or 4 in the breathing rate index and/or breathing quality and/or response to stimulation and/or level of consciousness will be observed.

#### Tolerability

The tolerability of test articles was assessed in healthy mice. Animals were treated by oral gavage (see below) at 10 mL/kg (ca. volume of 0.2 mL) twice daily (eight hours apart) at the different dosages (which represents the total daily dosage). The animals were be given the drugs for four days, and monitoring was continued 24 hours after the last treatment.

The following dosages were employed representing a 1:1 drug combinations: (i) 1.5 mg/kg Arainosine and 1 mg/kg Theobromine, (ii) 4.5 mg/kg Arainosine and 3 mg/kg Theobromine, (iii) 15 mg/kg Arainosine and 10 mg/kg Theobromine, (iv) 45 mg/kg Arainosine and 30 mg/kg Theobromine and, (v) 150 mg/kg Arainosine and 100 mg/kg Theobromine. All drugs were dissolved in Emulphor-EL-620. Each group contained three animals.

#### Anti-viral activity

Anti-viral efficacy studies were conducted on mice which were divided into seven groups, each containing six to eight mice. Each group of mice was intraperitoneally anesthetized (80 mg/mL Ketamine + 10 mg/kg Xylazine IP) and infected at the start of day one with 10,000 viruses per animal by intranasal administration of 40 μL. The inoculum size was determined as one that induced appreciable disease symptoms without reaching lethality. Subsequently each group was treated by oral gavage (10 mL/kg) BID, according to the treatment specified below, except for day one in which the animals received only a single treatment of half a daily dosage. The seven groups were treated as follows: (i) 1% Emulphor-EL-620 (vehicle control), (ii) Oseltamivir 20 mg/kg, (iii) 1.5 mg/kg Arainosine and 1 mg/kg Theobromine, (iv) 4.5 mg/kg Arainosine and 3 mg/kg Theobromine, (v) 15 mg/kg Arainosine and 10 mg/kg Theobromine, (vi) 45 mg/kg Arainosine and 30 mg/kg Theobromine and, (vii) 150 mg/kg Arainosine and 100 mg/kg Theobromine. All the combined drugs in desired concentrations were dissolved in an aqueous solution of 1% Emulphor-EL-620 while Oseltamivir was dissolved in water.

The experiment’s duration was five days during which all clinical signs were recorded including weight. After five days the animals were sacrificed and lungs were collected for further RNA quantification by RT-qPCR.

### Chemicals

The Isopropyl-*β*-D-1-thiogalactopyranoside was purchased from Biochemika-Fluka (Buchs; Switzerland). Xanthine and 3-methyl xanthine were purchased from Acros Organics (Antwerpen, Belgium). 1-methyl xanthine and Paraxanthine were purchased from ChemScene (Monmouth Junction, NJ, USA). Arainosine was purchased from BOC Sciences (Shirley, NY, USA). Enprofylline and 7-methyl xanthine were purchased from Glentham Life Sciences (Corsham, United Kingdom) and Alfa Aesar (Ward Hill, MA, US), respectively. All other chemicals were purchased from Sigma-Aldrich laboratories.

## Results

### Channel activity in bacteria

The channel activity of M2 was examined in bacteria, whereby conductivity that results from the protein changes the host’s phenotype. Specifically, three bacteria-based assays, negative, positive, and pHlux assays, were used to ensure reliability and minimize false results.

In the negative assay (Fig. 1a), increased protein expression upon raising Isopropyl-*β*-D-1-thiogalactopyranoside levles causes retardation of bacterial growth due to excessive membrane permeabilization (23). For example, at an Isopropyl-*β*-D-1-thiogalactopyranoside concentration of 45 μM, the bacterial growth is roughly halved compared to bacterial cells without an inducer.

In the positive assay, which is reciprocal to the negative assay, K^+^-uptake-deficient bacteria are used because they can only grow in a high potassium medium (19). However, these bacteria can also survive in low K^+^ medium when they express a K^+^ channel, which leads to renewed bacterial growth. Fig. 1b shows that appreciable bacterial growth is observed upon induction of protein expression by Isopropyl-*β*-D-1-thiogalactopyranoside and potassium supplement. Note that lower Isopropyl-*β*-D-1-thiogalactopyranoside concentrations are used in this assay since excessive permeabilization ensues at higher protein expression levels, and the benefit of potassium transport is nullified (16).

The final test is the pHlux assay (20) where a H^+^ influx into the bacteria is measured by a pH-sensitive GFP (21). Upon injection of acid into the medium, a fluorescence change will be observed due to cytoplasmic acidification that results from H^+^ influx mediated by the protein. Furthermore, different protein expression levels that result from different Isopropyl-*β*-D-1-thiogalactopyranoside concentrations will yield different acidification rates. As seen in Fig. 1c, significant acidification is observed when the inducer concentration increases to 50 μM. However, lower acidification is observed at inducer concentrations beyond 50 μM, most likely due to the detrimental impact of the protein on bacterial growth, as observed in the negative assay (Fig. 1a).

In conclusion, all three assays indicate appreciable conductivity by the viral protein, manifested by a change in bacterial phenotype. Consequently, reversing the phenotype can be used in drug screening, as expounded below.

### Drug screening

We employed the above bacteria-based assays to search for compounds that can inhibit (*i*.*e*., block) the activity of the Influenza M2 channel. In particular, we used the negative assay as the primary screening tool since, in this instance, successful blockers will be identified due to their ability to enhance bacterial growth. Hence, the negative assay presents a stringent test requiring bacteria to grow faster than otherwise, thereby filtering out any compounds that are toxic to the host. In contrast, in the positive assay, successful compounds will be deleterious to growth, and in the pHlux assay, they will abrogate fluorescence change. Rimantadine was used as a positive control for all assays, while no drug treatment served as the negative control. In each of the three assays, an optimal protein expression level was selected to maximize the dynamic range of blocker activity. Finally, we employed a repurposed drug library numbering 2,839 compounds. Such a library represents a set of compounds with known toxicity and can be used as a starting point for further chemical exploration and subsequent examination of anti-viral effect in mammalian cells and *in vivo* experiments.

The above process led to seven compounds designated as hits against the aminoadamantane-sensitive channel (Fig. 2a): Amikacin, Asunaprevir, Fludarabine, Flunisolide, Kasugamycin, Ravuconazol, and Theobromine. The same screening process was then employed on the M2 channel with the S31N aminoadamantane-resistant mutation (17) yielding eight hits of which five were active against the aminoadamantane-sensitive channel as well (Fig. 2b): Alvimopan, Amikacin, Asunaprevir, Emamectin, Fludarabine, Levamlodipine, Ravuconazol, and Theobromine. ote that as expected, Rimantadine was a potent blocker of the aminoadamantane-sensitive channel, but was entirely ineffective against the channel with the S31N mutation that is known to be aminoadamantane-resistant (17). Finally, all hits were also examined against their respective channel using the positive and pHlux assays with varying activity (Supporting Fig. 2).

**Fig 2.**
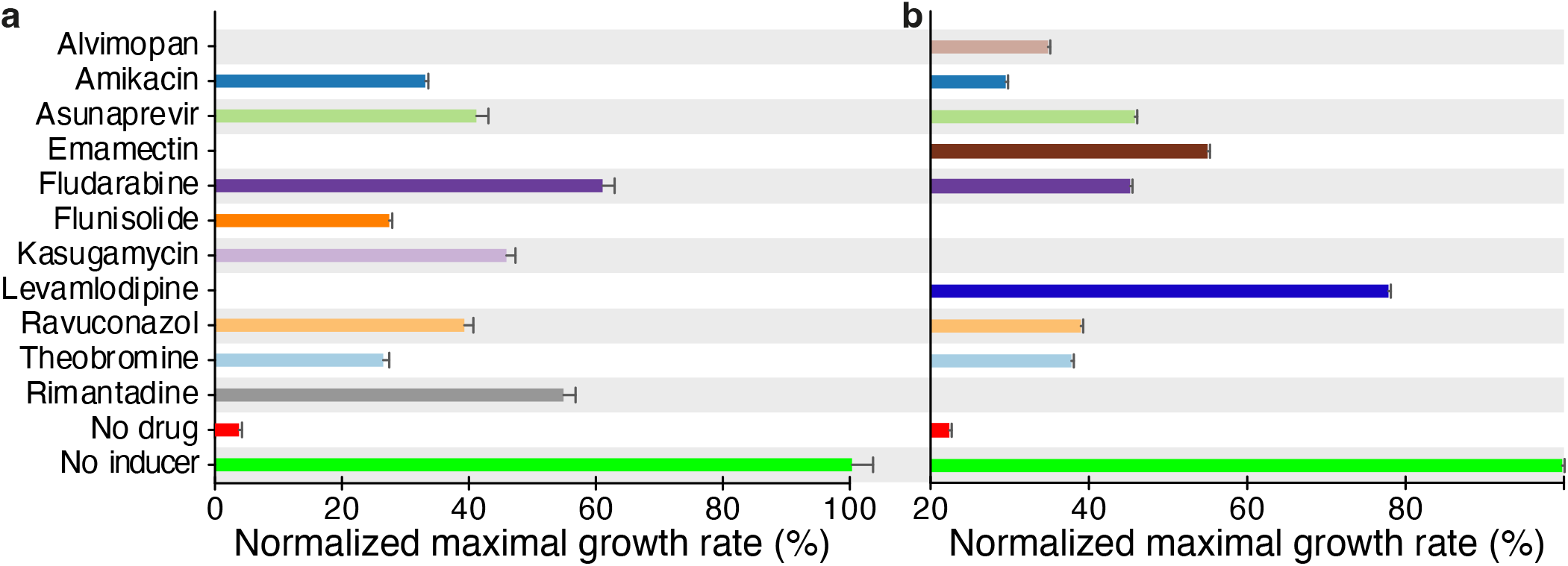
Screening results employing the negative assay against the aminoadamantane-sensitive M2 channel (a) and against a channel containing the S31N mutation (b) that renders the channel aminoadamantane resistant (17). Bacteria without the Isopropyl-*β*-D-1-thiogalactopyranoside inducer (green) and untreated bacteria (red) serve as controls. Effective blockers increase bacterial growth. Note that rimantadine exhibits potency against the aminoadamantane-sensitive channel (a) but, as expected, has no activity against a channel with the S31N mutation and is incapable of restoring bacterial growth (b).

#### *In-cellulo* anti-viral activity

We proceeded to examine the antiviral activity of the blockers we identified in the bacteria-based assays. To that end, the pandemic H1N1/09 swine flu strain was selected since it was the first virus designated by the World Health Organization as a public health emergency of international concern. In addition, the H1N1/09 virus contains the S31N mutation circulating in the vast majority of current viruses (24), rendering them resistant to aminoadamantanes (17). The assay was based on the ability of the hit compounds to reduce viral-induced cellular death in tissue culture. Oseltamivir and uninfected cells served as positive controls, while untreated cells as negative controls

#### Compound toxicity assessment

Prior to efficacy analysis, it as imperative to determine the inherent toxicity of the different hits. Hence, a tolerability test was performed to confirm the dose range up to which the hit concentration it is safe to use in the efficacy studies below. Specifically, we tested the cellular toxicity of each compound at 1, 3, 10, and 30 μM concentrations, by monitoring the cell viability after 48 hrs. As shown in Supporting Fig. 3, all compounds do not exhibit any toxicity up to 30 μM. The only exception is Levamlodipine, that exhibits increasing toxicity from 3 μM onwards.

**Fig 3.**
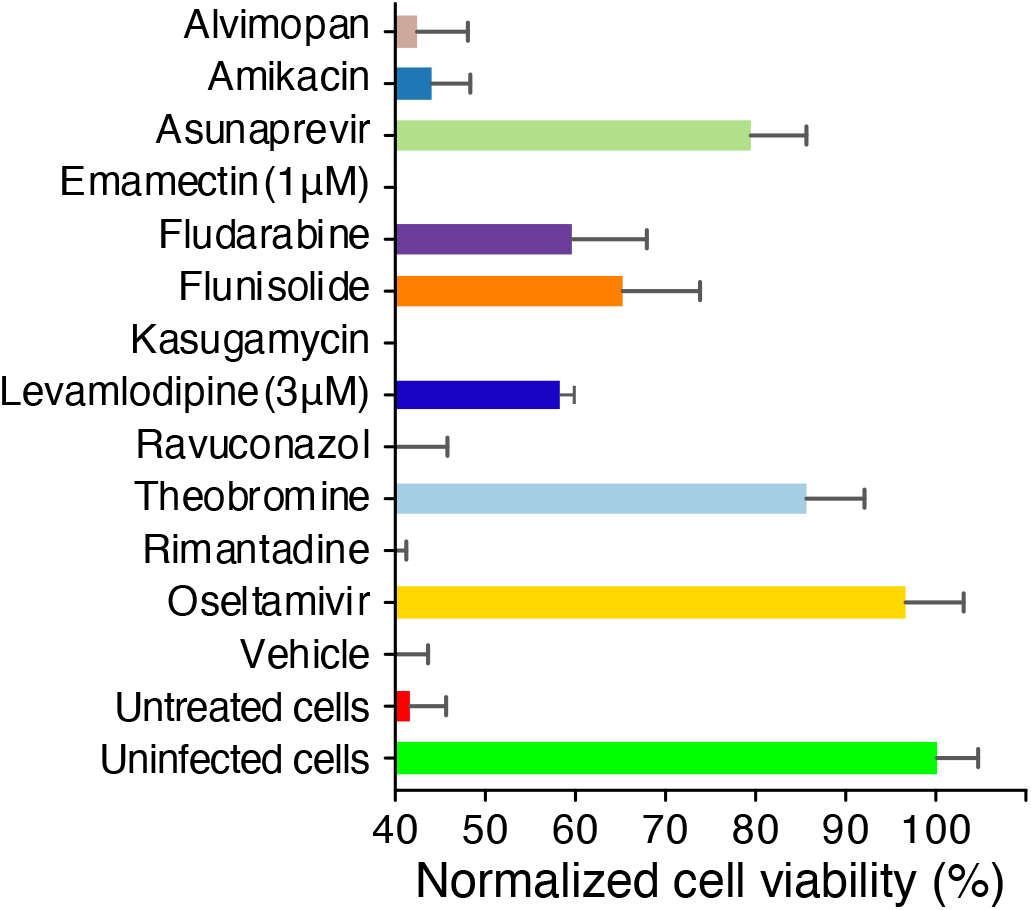
Anti-viral activity of M2 channel blockers. MDCK cells were infected with an H1N1 virus at an MOI of 0.3 and their viability was monitored by MTS after 48 hrs. Effects of different drugs at a concentration of 10 μM in 0.1% DMSO are listed. Emamectin and Levamlodipine were tested at 1 and 3 μM concentration, respectively, due to their toxicity. Results are normalized relative to uninfected cells.

#### Anti-viral efficacy of hit compounds

Having examined the toxicity of the different hits, we could examine their anti-viral activity *in-cellulo*. With an A/Wisconsin/629-D02452/2009 (H1N1)pdm09 virus with an S31N M2 channel, we tested the ability of compounds to reduce viral-induced death. The compounds were tested at 10 μM (or lower if toxicity studies did not enable such a high concentration), and the viability of the cells was examined two days after infection.

The results shown in Fig. 3 demonstrate that at an MOI of 0.3, more than 55% of the cells are non-viable after two days due to viral infection. However, several of the hits obtained from the bacterial screening were found to increase cell viability considerably. In particular, Asunaprevir, Fludarabine, Flunisolide, Levamlodipine, and Theobromine exhibited appreciable activity in negating virus-induced cellular death. Oseltamivir, a neuraminidase inhibitor, serves as a positive control and can almost entirely prevent viral-induced death.

#### Activity of structurally similar compounds

All five compounds that exhibited appreciable *in cellulo* activity at 10 μM were subjected to a search for structurally similar compounds. In doing so, we could fully utilize the repurposed drug library and uncover additional compounds that may not have scored positively in the bacteria-based assay but could still possess anti-viral activity.

A search for compounds similar to Fludarabine yielded Vidarabine, while the search for Theobromine yielded Theophylline, and Asunaprevir yielded Grazoprevir. Other search outcomes yielded compounds that did not exhibit any antiviral activity. We then conducted a dose-response analysis for all the above-mentioned active compounds, including both the parent and similar compounds. Two commercially available anti-influenza drugs, Oseltamivir and Favipiravir (viral RNA-dependent RNA polymerase inhibitor) were used as positive controls.

Results shown in Fig. 4 indicate that the following compounds exhibit activity with a well-defined dose-response characteristic: Asunaprevir (*Ks* = 60 nM, Bmax = 71%), Fludarabine (*Ks* = 32 nM, Bmax = 50%), Flunisolide (*Ks* = 47 nM, Bmax = 47%), Grazoprevir (*Ks* = 41 nM, Bmax = 64%), Theobromine (*Ks* = 387 nM, Bmax = 50%), and Vidarabine (*Ks* = 411 nM, Bmax = 66%). However, individually, none of the compounds was as efficacious as Oseltamivir with a *Ks* of 17 nM and a Bmax of 101%.

**Fig 4.**
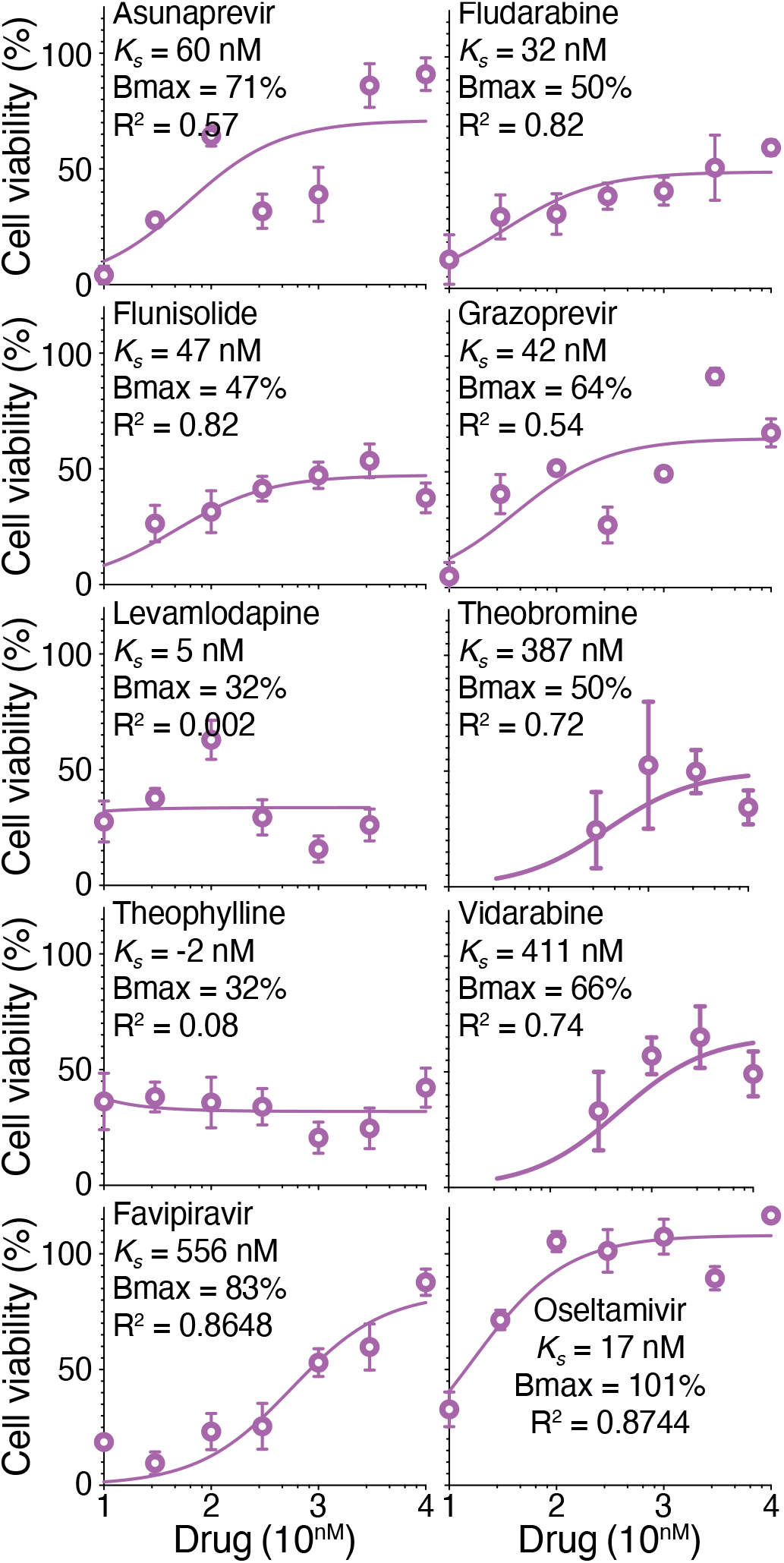
*In cellulo* anti-viral dose-response studies of active drugs. MDCK cells were infected with an H1N1 virus at an MOI of 0.3 and their viability was monitored by MTS after 48 hrs. Results are normalized relative to uninfected cells and untreated cells. *K*_*s*_, Bmax, and corresponding R^2^ values are shown for each drug in the inset.

#### Drug combination studies

To examine any beneficial antiviral activity of multiple drugs we studied the effect of drug combination at 0.1 μM as starting point. As shown in Fig. 5, few combinations showed pronounced synergism compared to their individual components. In particular, the combination of Theobromine and Vidarabine was able to completely protect cells from virus-induced cellular death, while individually, no activity was observed. The combination of Vidarabine with Grazoprevir was also potent, but in this instance, Grazoprevir on its own restores viability to 56% compared to untreated cells. All other combinations exhibited additivity, or no added effect.

**Fig 5.**
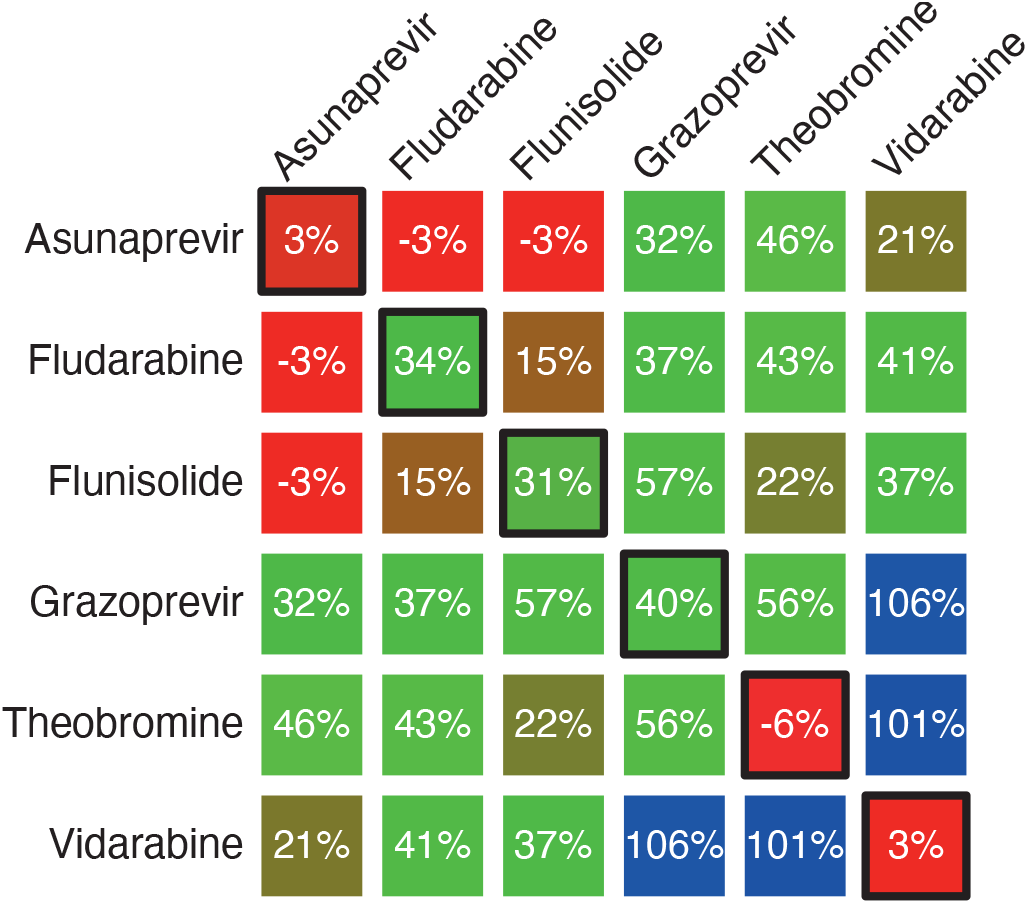
Combination anti-viral *in cellulo* studies of Asunaprevir, Fludarabine, Flunisolide Grazoprevir, Theobromine, and Vidarabine. MDCK cells were infected with the H1N1 virus at an MOI of 0.3 and their viability was monitored by MTS after 48 hrs. All compounds were at 0.1 μM. Diagonal elements (boxed) represent activity of the individual drugs. Results are normalized relative to uninfected cells and untreated cells and represent the average of at least three measurements.

Having identified a remarkable synergism between Theobromine and Vidarabine, we sought to characterize it further. In Fig. 6, we present the results of a combination study at concentrations from 10–300 nM. Once again, stark synergism is observed, but this time even at lower concentrations, whereby at an equal molar ratio of 30 nM, near complete protection from virus-induced cellular death is observed. For comparison, two approved anti-influenza drugs at 30 nM, Oseltamivir and Favipiravir, are able to provide only 71% and 10% protection, respectively.

**Fig 6.**
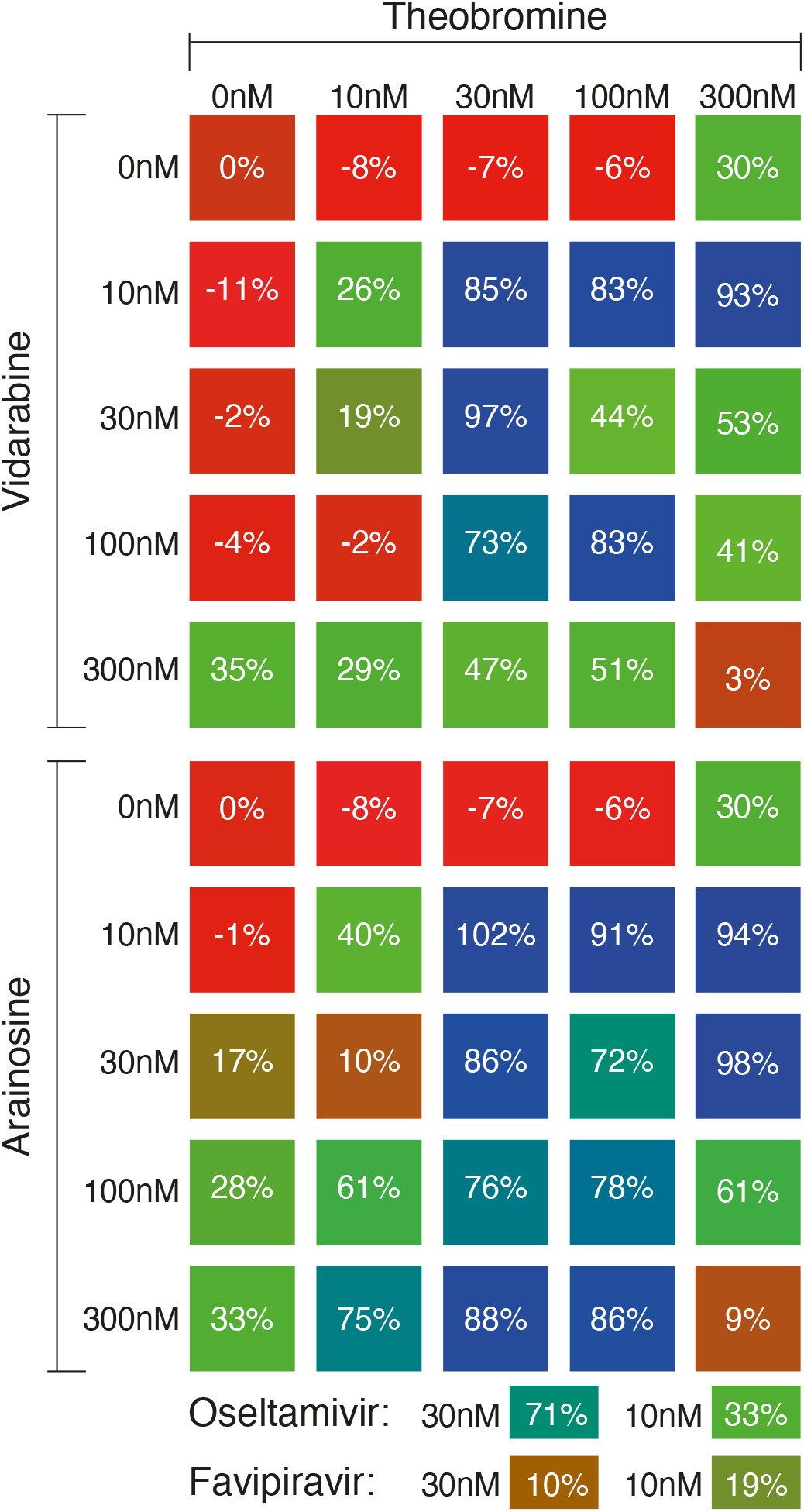
Synergy analyses of anti-viral *in cellulo* activity between Theobromine and Vidarabine and between Theobromine and Araionsine. The anti-viral activity of two approved drugs at 30 nM and 10 nM are shown at the bottom for comparison. MDCK cells were infected with H1N1 at an MOI of 0.3 and their viability was monitored by MTS after 48 hrs. Results are normalized relative to uninfected cells and untreated cells and represent the average of at least three measurements.

Interestingly, anti-viral synergism seems to decrease at higher concentrations. For example, at an equal ratio of 30 nM, both drugs exhibit 97% protection but the activity wanes to 83% and 3% at 100 nM and 300 nM, respectively.

Vidarabine is known to rapidly deaminate to Arainosine by adenosine deaminase (25). Therefore, we decided to examine the activity of Arainosine as well, individually and in combination with Theobromine. As see in Fig. 6, individually, Arainosine is perhaps slightly more effective than Vidarabine, but both compounds exhibit ca. 34% activity at 0.3 μM, whereas Oseltamivir is 100% effective at that concentration. However, once again, dramatic synergism is observed, where complete protection from virus-induced death is observed at 10 nM of Arainosine and 30 nM of Theobromine. These results indicate that the active form of Vidarabine is its deaminated metabolite - Arainosine. Finally, a decrease in activity at higher concentrations is observed once again, but to a lesser extent.

#### Activity against aminoadamantane-sensitive virus

So far, the remarkable synergism of Theobromine and Arainosine has been demonstrated against an aminoadamantane-resistant viral strain. This pandemic 09 H1N1 virus contains the S31N mutation in its M2 channel, rendering it aminoadamantane-resistant (17), as shown experimentally in Fig. 3. Therefore, we repeated the experiments on an H5N1, highly pathogenic avian Influenza virus (Israel/975/2023) that, according to the sequence of its M2 channel, should be aminoadamantane-sensitive (Supporting Fig. 1). Results shown in Fig. 7 indeed demonstrate that the virus is aminoadamantane-sensitive, whereby Rimantadine is a potent inhibitor of infectivity, while against the H1N1 virus, it showed no activity whatsoever (Fig. 3). Furthermore, the *Ks* measured against the virus (15.3 nM) is indistinguishable from previous results (13±4 nM) obtained in a bacteria-based assay (23).

**Fig 7.**
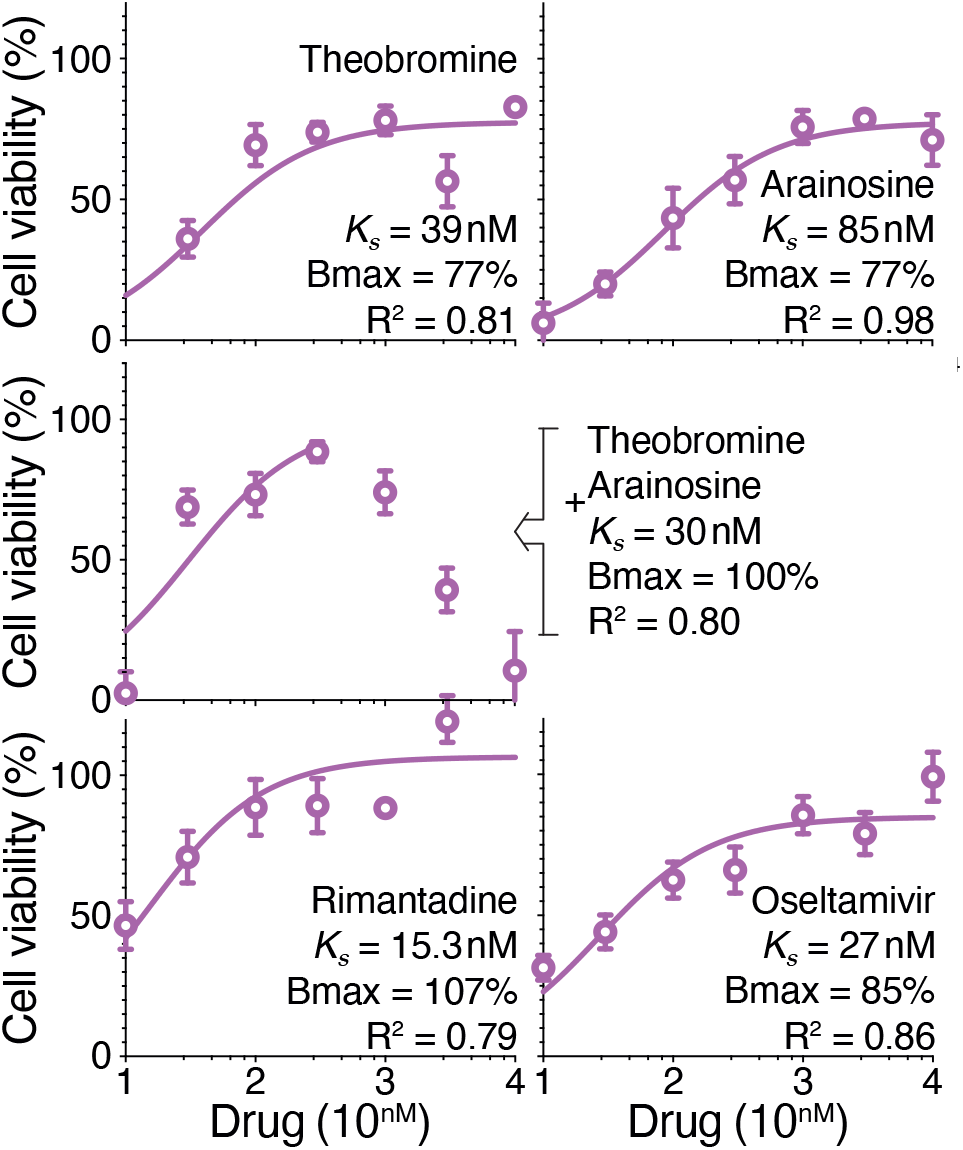
*In cellulo* anti-H5N1 dose-response studies of active drugs. MDCK cells were infected with the H5N1 virus at an MOI of 0.3 and their viability was monitored by MTS after 48 hrs. *K*_*s*_, Bmax, and corresponding R^2^ values are shown for each drug within the inset. Note that for the Theobromine and Arainosine drug combination, equal concentrations were used and the fit employed data until 0.3 μM. Results are normalized relative to uninfected cells and untreated cells.

Turning to the drug duo, both Theobromine and Arainosine individually exhibited activity against the aminoadamantane-sensitive avian Influenza (Fig. 7). Yet, the activity of the individual compounds is significantly better against the aminoadamantane-sensitive strain relative to the activity against the aminoadamantane-resistant strain, as summarized in Table 1. For example, Theobromine exhibited an affinity better by an order of magnitude alongside higher maximal activity. However, combining the two drugs against the aminoadamantane-sensitive strain leads only to mild additivity rather than stark synergism, which is observed against the aminoadamantane-resistant strain. Specifically, when both drugs are combined at 30 nM, similar protection from viral-induced death is observed against the aminoadamantane-sensitive and aminoadamantane-resistant strains (69% and 86%, respectively).

**Table 1.** Anti-viral parameters of selected compounds against aminoadamantane-resistant and aminoadamantane-sensitive strains. The data are tabulated from Fig. 4 and Fig. 7, respectively.

| Strain | Aminoadamantane sensitivity | Theobromine |  | Arainosine |  | Oseltamivir |  | Rimantadine |  |
| --- | --- | --- | --- | --- | --- | --- | --- | --- | --- |
| | | $K_s$ | Bmax | $K_s$ | Bmax | $K_s$ | Bmax | $K_s$ | Bmax |
| H1N1 | Resistant | 400 nM | 50% | 400 nM | 66% | 17 nM | 101% | No activity |  |
| H5N1 | Sensitive | 39 nM | 77% | 85 nM | 77% | 27 nM | 85% | 15 nM | 107% |

#### Inhibition site

In order to ascertain the target of the Theobromine and Arainosine drug duo, we examined its binding properties to the M2 protein. Previously, we have shown that adamantanol is a competitive inhibitor of aminoadamantanes (26). Adamantanol shares the adamantane moiety that enables it to bind the M2 channel but lacks a positive charge that repels H^+^ permeation. Consequently, we tested if adamantanol can inhibit the activity of Theobromine, Arainosine, and their combination in the bacteria-based assay and *in cellulo*.

As seen in Fig. 8, amantadine can effectively relieve viral-induced cellular death caused by the H5N1 virus. It does so by blocking the viral channel, which is essential to its infectivity cycle (13). However, addition of excess of adamantanol (5 μM) abrogates the utility of amantadine (300 nM) almost entirely because it is its competitive inhibitor (26). Similarly, the activity of amantadine in the bacteria-based assays is nullified by adamantanol: In the negative assay, the ability of amantadine to relieve the growth inhibition due to excessive membrane permeabilization of the channel is negated by adamantanol (Supporting Fig. 5a). In the pHlux assay, amantadine blocks the H^+^ through the channel, but adamantanol counteracts this activity (Supporting Fig. 5b).

**Fig 8.**
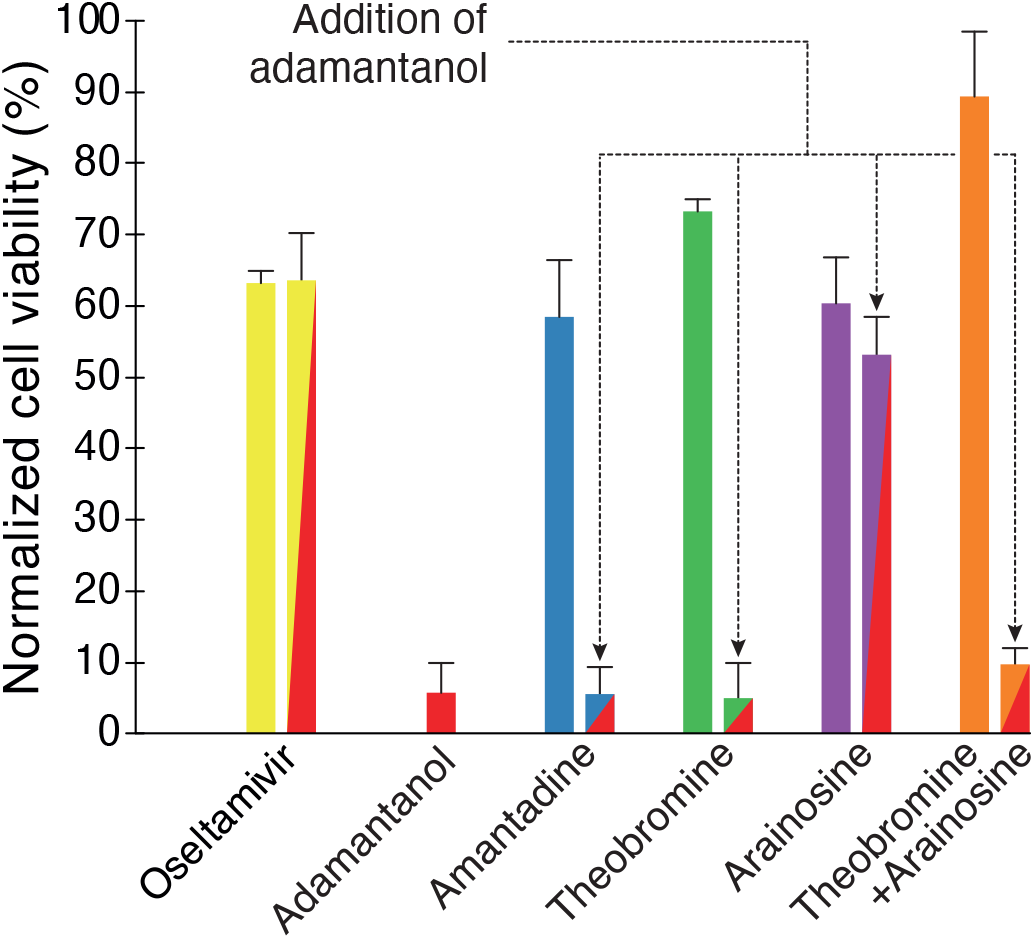
*In cellulo* binding mechanism analyses of Theobromine and Arainosine. MDCK cells were infected with the H5N1 virus at an MOI of 0.3, and their viability was monitored by MTS after 48 hrs. All drugs are given at 300 nM except adamantanol, which, on its own or in combination with other drugs, is administered at 5 μM. Results are normalized relative to uninfected cells and untreated cells.

Repeating the above examination of adamantanol’s impact on the anti-viral activity of Theobromine, Arainosine, and their combination reveals a similar picture (Fig. 8). Theobromine at 300 nM can provide cells with 73% protection from viral-induced death. However, adding adamantanol reduces the protection to a mere 5%. In Arainosine, a much smaller reduction is observed: The drug provides cells with 60% protection, which drops to 53% upon adding adamantanol. However, the activity of adamantanol is particularly pronounced in the Theobromine-Arainosine combination. In this instance, the drug duo provides cells with 89% protection from viral-induced death that drops to 10% upon adamantanol addition. Additionally, as expected, adamantanol has no protective effect against the virus, nor can it vitiate the benefit of Oseltamivir, which targets the viral neuraminidase. Finally, these results are once more mirrored in the bacteria-based assays shown in Supporting Fig. 5.

#### Structural activity relationship study

Given the remarkable synergism between Theobromine, a xanthine analog, and Vidarabine or Arainosine, both nucleoside analogs (Fig. 6), we sought to explore its specificity by examining highly similar analogs. In particular, we determined the anti-viral activity of individual xanthine and nucleoside analogs and combinations thereof.

Theobromine is a xanthine-group member and found to have available structural analogues namely, Caffeine, Theophylline, Enprofylline, Paraxanthine, 7-Methylxanthine, Xanthine, 1-Methylxanthine, and 3-Methylxanthine (Fig. 9a). Among them, 7-Methylxanthine shows higher activity than Theobromine at 10 μM concentration. Apart from that, Xanthine and 3-methylxanthine show comparable activity as Theobromine at a higher concentration range (0.3–10 μM).

**Fig 9.**
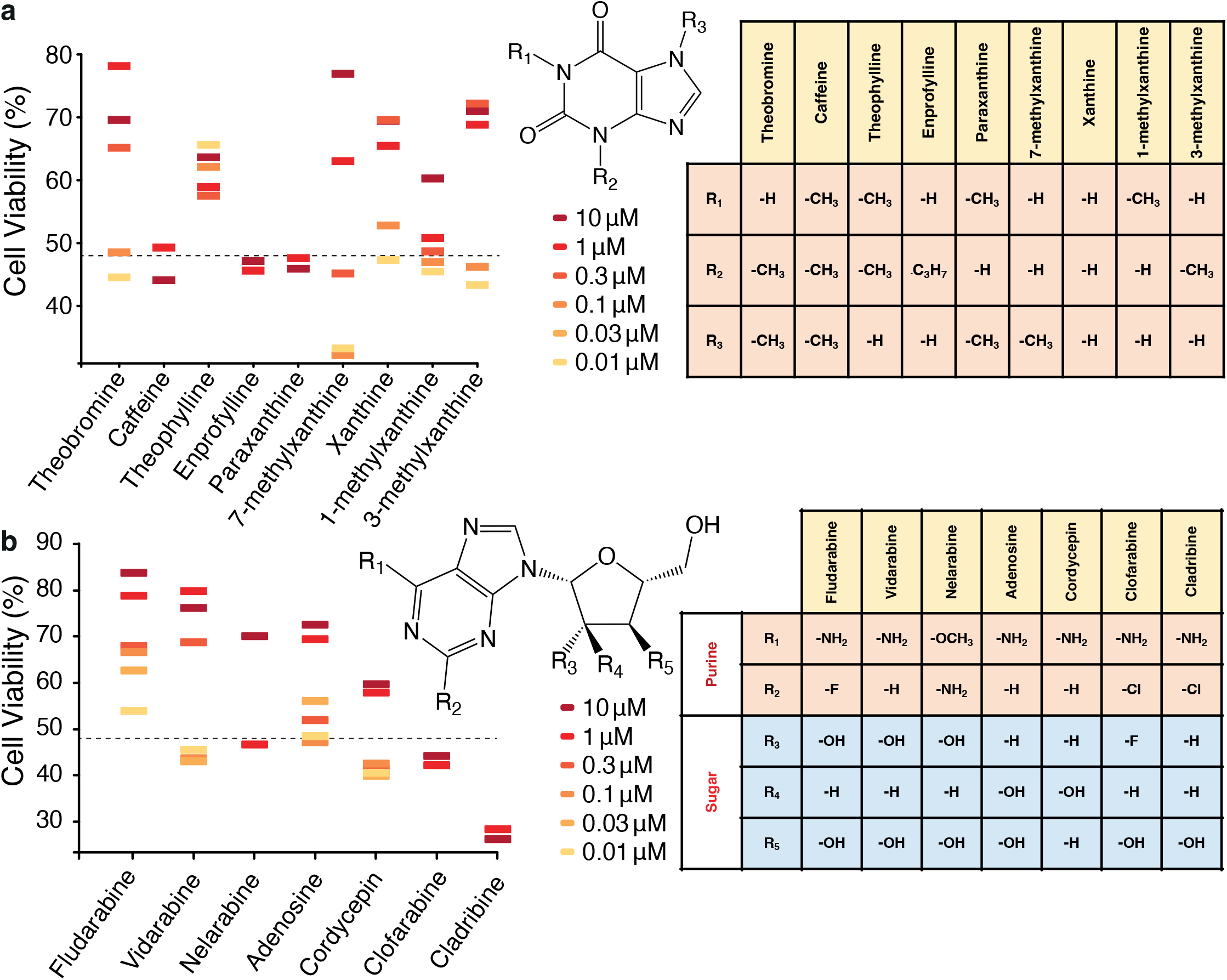
Structural activity relationship study of Theobromine (a) and Vidarabine (b). MDCK cells were infected with the H1N1 virus at an MOI of 0.3 and their viability was monitored by MTS after 48 hrs. Dashed lines indicate viability of untreated cells, while uninfected cells are set to 100%.

Vidarabine is a nucleoside analog of adenosine in which the ribose is epimerized to D-arabinose (27). Arainosine is the main metabolic byproduct of Vidarabine and is obtained by deamination (25). Therefore, we sought to compare the anti-viral activity of structurally similar analogs like Fludarabine, Nelarabine, Cordycepin, Clofarabine, Cladribine, and Adenosine.

Fig. 9b shows the activity of each drug at different concentrations along with the generalized structure of the nucleoside (inset and table). Fludarabine exhibits the highest activity, followed by Vidarabine, Adenosine, and Nelarabine. Other compounds have insignificant positive effects on cell viability.

We next proceeded to examine potential synergies between all of the nucleoside analogs and Xanthine analogs. As seen in Fig. 10, the only significant synergistic combination is between Theobromine and Vidarabine/Arainosine. Interestingly, even though Fludarabine was appreciably more potent than Vidarabine/Arainosine individually, it did not exhibit any synergism in combination with Theobromine (or any other Xanthine derivative).

**Fig 10.**
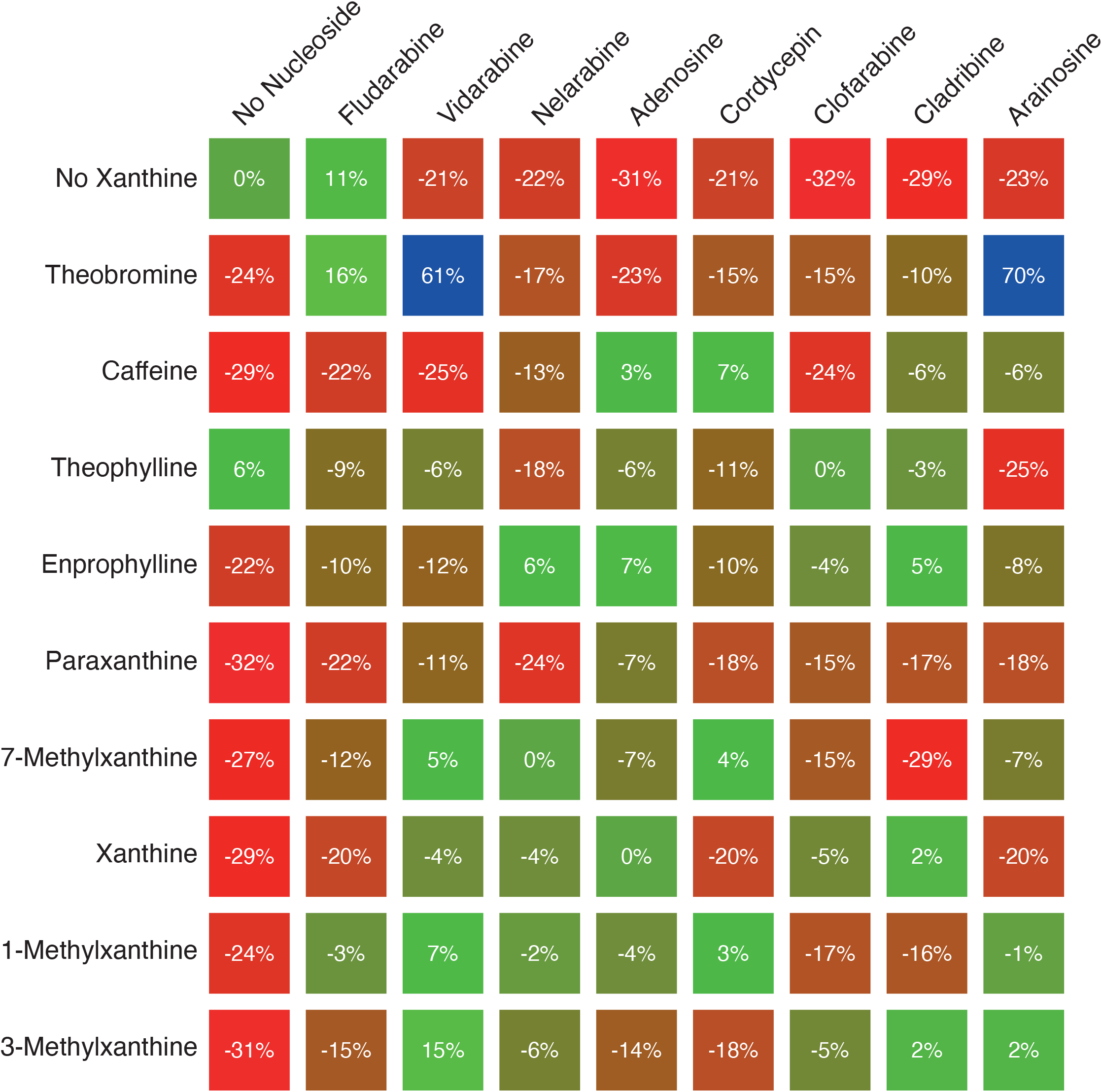
Combination anti-viral *in cellulo* studies of nucleoside analogs and Xanthine analogs. MDCK cells were infected with the H1N1 virus at an MOI of 0.3 and their viability was monitored by MTS after 48 hrs. Results are normalized relative to uninfected cells and untreated cells and represent the average of at least three measurements. All compounds were at at 100 nM.

#### *In-Vivo* study

After demonstrating the marked synergism between Theobromine and Arainosine, we examined whether the drug duo exhibits antiviral activity in animals. Initially, we established the drug duo’s safety in mice. While the individual drugs are known to be very tolerable, the impact of their combination has not been established. In brief, no adverse effects were observed in all dosages tested, up to and including 100 mg/kg of Theobromine and 150 mg/kg of Arainosine (data not shown).

To that end we infected mice with the Influenza strain A/Wisconsin/629-D02452/2009 (H1N1) pdm09, and monitored the amount of viral RNA in the lungs after five days. During the course of infection, the mice were treated with a the Theobromine-Arainosine drug duo at five different dosages. alongside Oseltamivir that served as a positive control, and a placebo that served as a negative control. All treatments were administered twice daily by oral gavage.

From the results shown in Fig. 11b, viral RNA load quantification clearly showed a reduction compared to placebo of the following drug combinations: 1 mg/kg Theobromine and 1.5 mg/kg and 3 mg/kg Theobromine and 4.5 mg/kg Arainosine (75% and 77% reduction, respectively). Oseltamivir at 20 mg/kg and 10 mg/kg of Theobromine and 15 mg/kg Arainosine showed a reduction in viral load to a lesser extent which was not statistically significant relative to the placebo (57% and 53%, respectively). At higher concentrations of 30 and 100 mg/kg Theobromine combined with 45 and 150 mg/kg Arainosine, respectively, the percentage of viral load reduction decreased.

**Fig 11.**
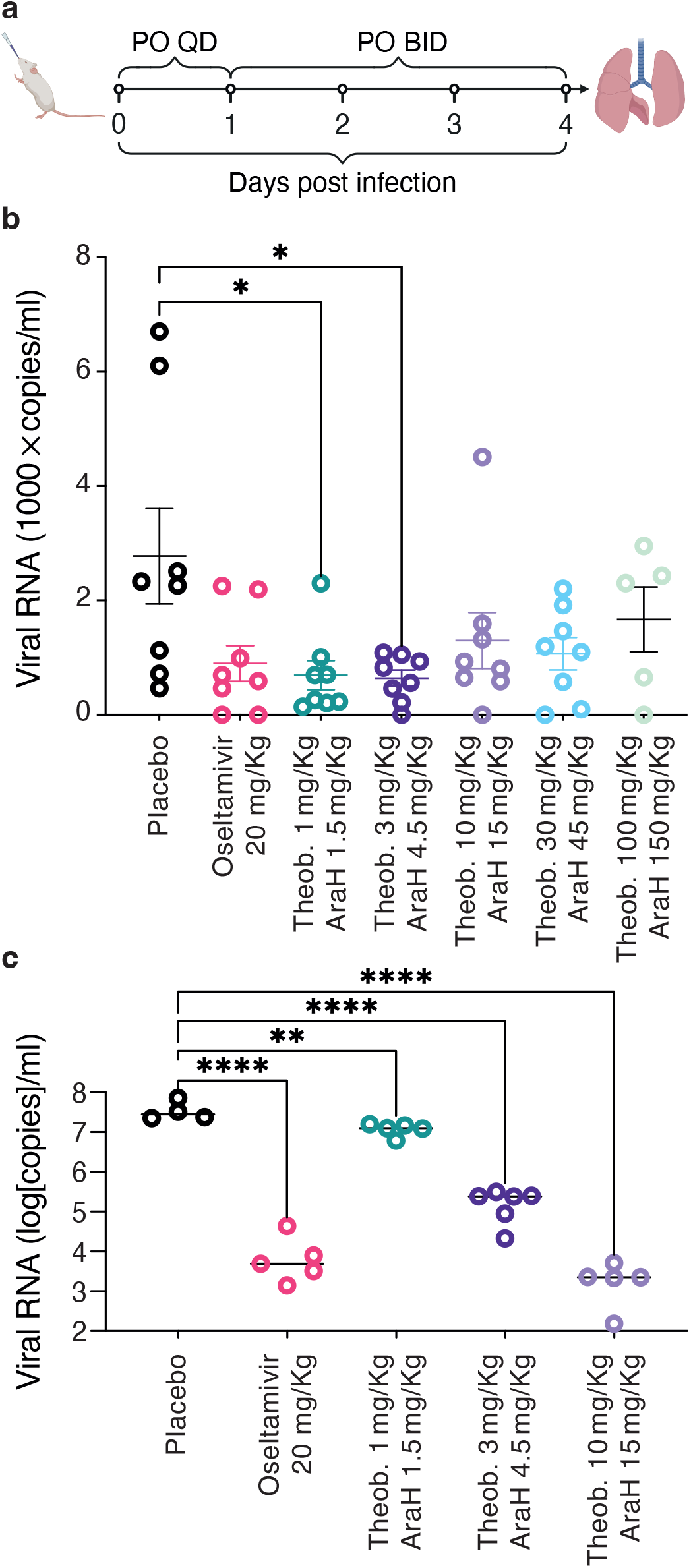
*In vivo* anti-viral activity. a. Scheme of experimental protocol. b. Mice were infected with the A/Wisconsin/629-D02452/2009 (H1N1)pdm09 strain and viral RNA in the lung was measured after five days c. Same as b, but mice were infected with the PR8 H1N1 strain.

The above influenza strain does not elicit pronounced disease symptoms in the mice. Therefore, we transitioned to the mouse adopted PR8 H1N1 strain that is particularly virulent (28). Note that results in tissue culture demonstrate indistinguishable susceptibility of the PR8 strain to Oseltamivir and the Theobromine-Arainosine duo (Supporting Fig. 4). As seen in Fig. 11c, pronounced differences are observed between all treatments and the placebo. Oseltamivir at 20 mg/kg, reduces the viral load by 3,077 fold, while the combination of 10 mg/kg Theobromine with 15 mg/kg Arainosine leads to a reduction of 15,717 fold. Lower concentration of Theobromine with Arainosine yield statistically significant reductions as well.

Since mice infected with the PR8 strain exhibited pronounced disease symptoms, we could compare the effects of the different treatments thereupon. In particular, we monitored the mice’s weight as an indicative metric for disease progression. As shown in Fig. 12, mice that were treated with the placebo exhibited severe symptoms, losing ca. 20% of their body weight in five days. Oseltamivir at 20 mg/Kg, as a positive control reduced the weight loss to less than 15%. However, the drug duo was significantly more effective once the dosage was raised to 3 mg/Kg Theobromine combined with 4.5 mg/Kg Arainosine. At a dosage of 10 mg/Kg Theobromine combined with 15 mg/Kg Arainosine the weight loss was reduced even further to only 6% after five days.

**Fig 12.**
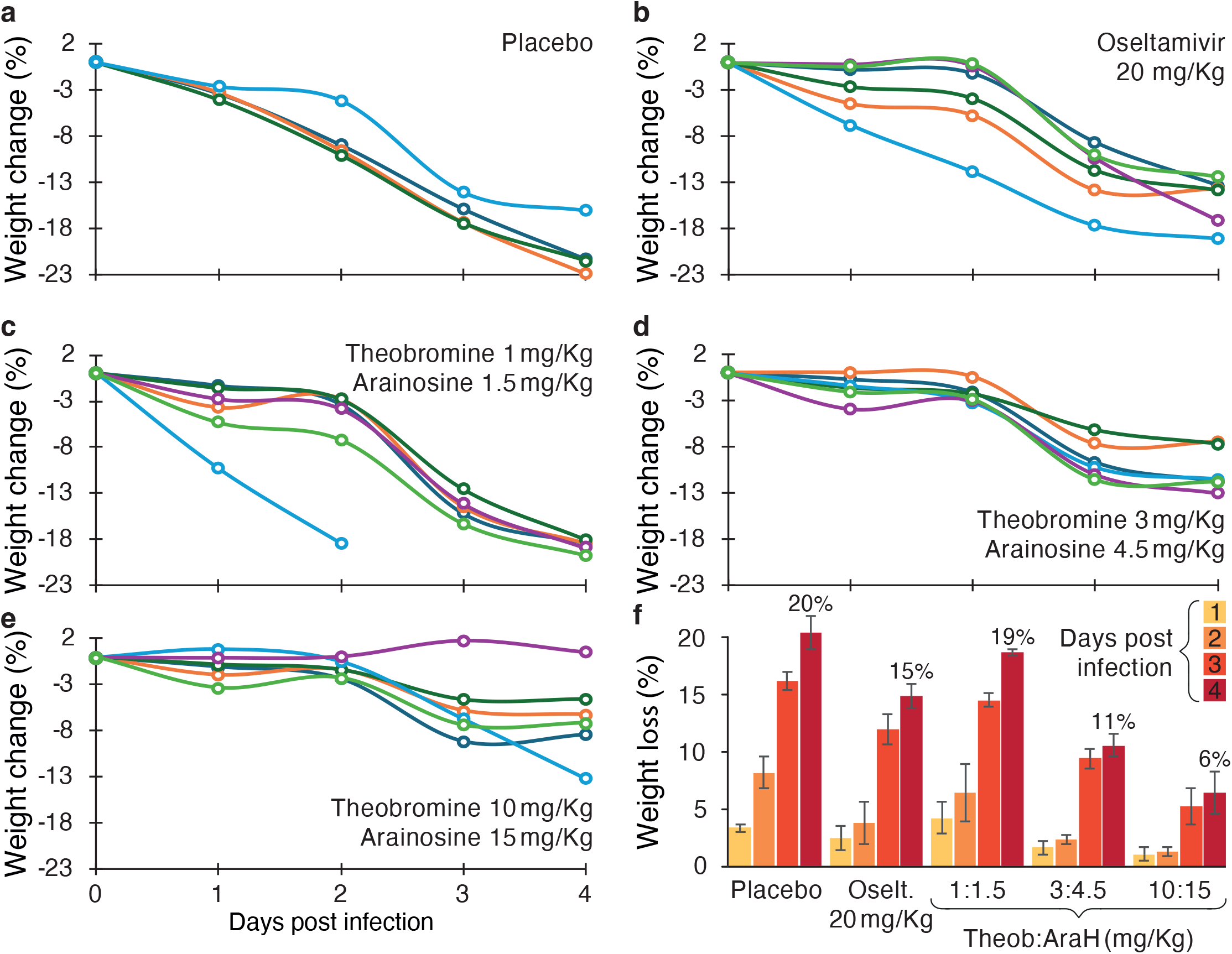
*In vivo* anti-viral activity. Relative weight changes of mice receiving different treatments, as indicated, during the course of the infection. a-e: Individual mice weight changes. f: Average data for comparative purposes whereby values of four days post infection are listed in the graph.

## Discussion

Influenza poses a severe health challenge, yet there is a dearth of curative anti-viral agents. Furthermore, the virus’s constant genetic changes continuously vitiate even the partial benefit of current treatments. Consequently, we have searched for potential anti-viral drugs for clinical application by attempting to identify new inhibitors of the virus’s viroporin.

Our approach, schematically shown in Fig. 13, involved several steps: (i) Identification of hits from a repurposed drug library with bacteria-based assays. (ii) Addition of similar compounds. (iii) Analysis of anti-viral activity in tissue culture. (iv) Combination studies to determine additivity and synergism. (v) Efficacy determination in animals. Finally, we focused on two of the most important strains that represent a pandemic threat: H1N1 swine flu and H5N1 avian flu. Moreover, we searched for agents that are active against both aminoadamantane-sensitive and aminoadamantane-resistant viral strains.

**Fig 13.**
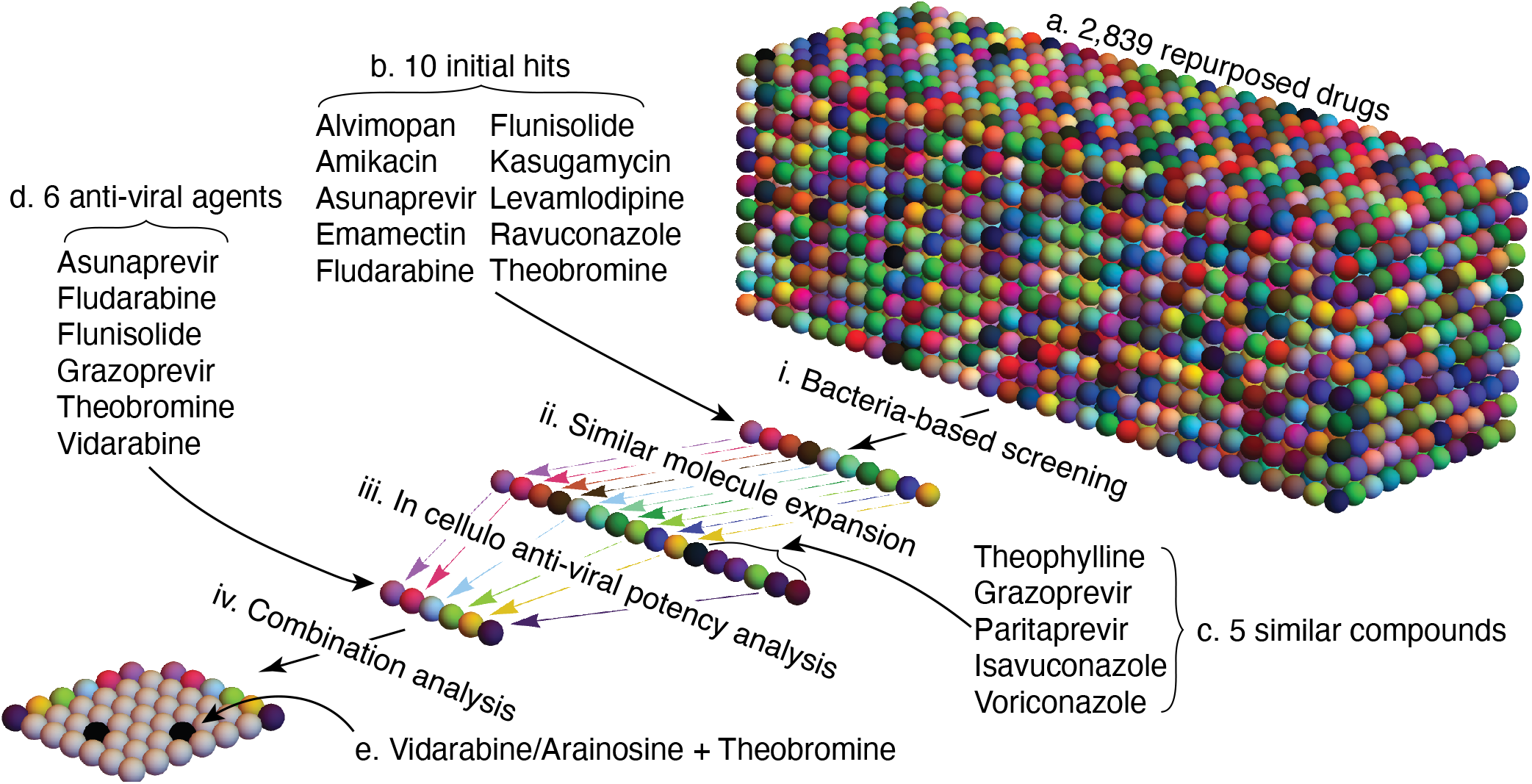
Systematic approach for anti-influenza agents yielding the Theobromine and Arainosine drug duo: i. A repurposed drug library of 2,839 compounds (a) was screened with bacteria-based assay to yield ten hits (b). ii. Five additional compounds (c) were found based on their similarity to the ten hits from the previous stage (b). iii. In cellulo testing demonstrated that six compounds (d) exhibit anti-viral activity out of the 15 expanded hits (b+c). iv. One pair of compounds exhibits remarkable anti-viral synergism from combination studies in tissue culture. Note that two combinations are shown due to the matrix’s symmetry.

Aminoadamantanes have long been known to possess potent anti-influenza activity (11) by targeting the M2 protein (12) through blockage of its channel activity (13). However, resistance emergence against aminoadamantanes is widespread, and most circulating strains contain the S31N mutation confers resistivity against the blockers (14). Therefore, our screening approach searched for blockers against the aminoadamantane-sensitive and aminoadamantane-resistant channels.

To examine the activity of the M2 channel and identify inhibitors thereof, we used bacteria-based assays. In such assays, the viral channel changes the host’s phenotype, affording all of the benefits of bacterial genetics. Such assays are particularly suited for viral channels, which, due to their small size, are routinely expressed in an active form in their bacterial host (20, 23, 29–34). As seen in Fig. 1, the M2 protein scored positively in all three bacteria-based assays, facilitating drug screening by selecting compounds that reverse the host’s channel-driven phenotype.

We screened 2,839 approved and advanced-stage drugs using three bacteria-based assays. A repurposed drug library, such as the one used, represents a subset of chemicals whose toxicity has been established and can serve as a starting point for further exploration. The bioassays yielded several compounds as hits that showed activity in both positive and negative assays. The hits were further checked by a fluorescence-based assay that monitors H^+^ flux to confirm channel-blocking activity.

Four successful compounds, Asunaprevir, Amikacin, Ravuconazole, and Fludarabine, exhibited potential blocking activity (Fig. 2a). They were active against an aminoadamantane-sensitive channel and showed activity when checked on mutated S31N M2 channel known to be aminoadamantane-resistant (17). Apart from these, Theobromine and Flunisolide showed promising activity in the negative and fluorescence-based assays. The three bacteria-based assays monitor similar, albeit not identical, properties, which could be why compounds may exhibit different activities in the different assays. In addition, another set of experiments was performed to screen all the drugs on the M2 S31N aminoadamantane-resistant channel; several compounds showing activity on negative assays were also selected for further studies. These are Alvimopan, Emamectin, and Levamlodipine (Fig. 2b).

All the drugs selected from the bacteria-based screening effort were then subjected to *in cellulo* anti-viral activity assays. This test examined compounds in tissue culture by their ability to reduce viral-driven death. To increase the scope of the study, we also included similar compounds to the hit molecules obtained from bacterial assays. Among the selected compounds, it was observed that Asunaprevir, Fludarabine, Flunisolide, Levamlodipine, and Theobromine protect cells from viral-induced death at 10 μM concentration (3 μM in the case of Levamlodipine), albeit at levels that are lower than Oseltamivir (Fig. 3). Furthermore, since the H1N1 strain contains the S31N mutation that renders it aminoadamantane-resistant, it is of no surprise that Rimantadine exhibited no activity whatsoever. Finally, dose-response studies of the above active blockers, together with similar compounds, Vidarabine (similar to Fludarabine) and Grazoprevir (similar to Asunaprevir), yielded sub-micromolar *Ks* values of several hits (Fig. 4). However, individually, Oseltamivir is more active in its affinity and its maximal inhibition.

There is always a risk that compounds identified by their ability to inhibit a particular viral component in isolation inhibit the whole virus by an off-target route. In other words, while the compounds were selected based on their ability to block the M2 channel, they may exert their anti-viral activity through different mechanisms. However, the fact that several unrelated compounds retrieved from the bacteria-based selection for channel blockers exhibit anti-viral activity suggests a low probability of an off-target effect. Hence, it is doubtful that the compounds inhibit the virus by any mechanism other than by blocking its channel.

It is possible to rule out an off-target mechanism altogether by utilizing adamantanol. The drug is an analog of amantadine but lacks a positive charge and, consequently, cannot block the M2 channel (26). However, because it shares the adamantane group with aminoadamantanes, it can bind the channel and competitively inhibit their function (26). The competitive inhibition of adamantanol can also be seen when analyzing activity in tissue culture where it abrogates the anti-viral activity of amantadine almost entirely (Fig. 8). Encouragingly, adamantanol has the same neutralizing activity on Theobromine, Arainosine (to a lesser extent), and their combination (Fig. 8). Hence, it is challenging to conjure up a scenario in which the drugs inhibit the virus by any mechanism other than blocking its M2 channel. Finally, as expected, adamantanol has no anti-viral effect on its own, nor does it impact the activity of Oseltamivir since the drug works by an entirely different mechanism as a neuraminidase inhibitor (35).

The additive effect of more than one drug is another integral part of drug research and has been found to be effective in potentially curing several diseases (36). In the current study, we found more than one effective drug against the Influenza virus. This finding prompted us to check the combined effect of different drugs that showed a dose-response activity in the in cellulo studies. Interestingly, while most combinations exhibited mild additivity, if at all, the combinations of Grazoprevir + Vidarabine and Theobromine + Vidarabine showed activity beyond additivity when compared to the sum of their components (Fig. 5).

The combination of Theobromine and Vidarabine was particularly remarkable. Individually, Theobromine and Vidarabine can barely, if at all, protect tissue culture cells from viral onslaught: −6% and 3%, respectively. Yet astonishingly, together, they provide complete protection from viral-induced death. Consequently, we varied the concentrations of each component in a matrix format to further examine this synergistic interaction. The results depicted in Fig. 6 show that the drug combination exhibits exceptional potency even at lower concentrations. For example, both drugs combined at 30 nM provide near complete viability protection, while in comparison, on their own, they exhibit no activity. Oseltamivir, the leading drug on the market against influenza at 30 nM, exhibits only 71% protection. Furthermore, Vidarabine at 10 nM combined with Theobromine at concentrations equal to or above 30 nM exhibits activity higher than 83%. Oseltamivir, on the other hand, at 10 nM, can protect only a third of the cells from viral-induced death.

It is interesting to note that synergism is reduced at higher concentrations of both compounds. Mechanistically, Harris and co-workers have shown that sugars act as selective hydrotropes toward Caffeine (37). Therefore, it is possible to speculate that the sugar moiety of Vidarabine may similarly serve as a hydrotrope to Theobromine, which differs from Caffeine by a single methyl group.

Earlier evidence has found that the primary metabolic product of Vidarabine is Arainosine, which is produced through deamination by adenosine deaminase (38). Therefore, we tested whether Arainosine exhibits the same synergism with Theobromine. The results shown in Fig. 6b demonstrate that the combinations of these two drugs provide comparable results to the combination of Theobromine and Vidarabine. Hence, it is possible to conclude that Vidarabine is in fact a prodrug that is converted to Arainosine. Interestingly, this finding contradicts Vidarabine’s anti-cancer and anti-DNA virus activity, in which deamination renders the drug inactive (38). This result is perhaps unsurprising since the Influenza virus does not go through a “DNA phase” during its infectivity cycle. Hence, the activity of Arainosine on Influenza should employ a different mechanism (*i*.*e*., blocking the M2 channel), as the impact of adamantanol suggests. Finally, the synergism observed between Theobromine and Arainosine was diminished to a lesser extent at higher concentrations. Consequently, we chose to focus on Arainosine instead of Vidarabine from this point onwards.

All of the above studies were conducted on the H1N1 pandemic swine flu strain, which is aminoadamantane-resistant. Therefore, we decided to examine the activity of both compounds and their combination on an aminoadamantane-sensitive viral strain. For this purpose, we utilized the clinically important H5N1 avian influenza in which the particular variant is aminoadamantane-sensitive. In this instance, the results are very intriguing: An equimolar combination of Theobromine and Arainosine is similarly active against either the aminoadamantane-resistant or aminoadamantane-sensitive strains, providing near complete protection from viral infection at 30 nM (Fig. 7). To reiterate, the Theobromine-Arainosine drug duo is not only active against the important H1N1 swine flu and H5N1 avian flu strains, it is also active against aminoadamantane-resistant and aminoadamantane-sensitive strains alike.

Interestingly, the activity of the compounds, when used on their own, distinguishes the two strains. At 30 nM, no activity is observed for either drug against the aminoadamantane-sensitive H1N1 strain (Fig. 4 and Fig. 6). Yet, at the same concentration, both compounds exhibit appreciable but non-maximal activity against the aminoadamantane-resistant H5N1 strain (Fig. 7). Hence, either drug can inhibit (partially) the aminoadamantane-sensitive strain. At the same time, their combination is required for complete protection. In contrast, the individual drugs offer no benefit against the aminoadamantane-resistant variant, yet their combination provides full protection. Finally, it is interesting to note that at concentrations higher than 300 nM, a diminishing effect is observed in combination therapy for both viral strains, as discussed above.

Further insight into the stark synergy between Theobromine and Arainosine in the aminoadamantane-resistant strain, was obtained by exploring the activity of similar compounds, individually and in combination. We started with Theobromine, a Xanthine derivative, of which there are many commercially available analogs. In particular, we were able to examine every methyl derivative combination at positions R1, R2, and R3, as shown in Fig. 9a.

Comparison of Xanthine with 3- and 7-methylxanthine indicates that at higher concentrations, it is less active than both of its methylated derivatives. At lower concentrations, 3-methylxanthine is more efficacious than its 7-methyl analog. Yet, all three compounds are less effective than Theobromine. Therefore, it can be concluded that methylations at the 3 and 7 positions are necessary to yield optimal anti-viral effects.

1-methylxanthine, though, has an effect at a lower concentration range but does not show an effect at a higher concentrations. On the other hand, theophylline, which is methylated at the 1 and 3 positions is active to some extent but shows poor dose-dependent activity (Fig. 4). Notably, the 1,7-dimethylated analog, Paraxanthine, exhibits even lower activity. These observations further emphasize the role of methyl groups at the 3 and 7 positions, and any deviation from that leads to loss of activity. Moreover, the molecule completely loses all activity when all three nitrogen atoms are methylated, as in Caffeine. Finally, to check the activity of the alkyl group, we employed Enprofylline or 3-propylxanthine and found no effect when compared to its methyl analog 3-methylxanthine.

Arainosine or its metabolic precursor Vidarabine are nucleoside analogs in which a purine is n-linked to a pentofuranose. While not as numerous as the analogs of Theobromine, we examined several commercially available analogs for their antiviral activity in cell viability assays (Fig. 9a).

Fludarabine was the lead compound identified by the bacteria-based channel assays and in the *in cellulo* experiments. Therefore, compared to Vidarabine, a Fluorine at R2 in the purine ring (the only point of difference with Vidarabine) does not hamper the activity. On the other hand, substituting – NH2 at R1 with a – OCH3 group, as in Nelarabine, significantly reduces activity (only active at 10 μM).

Analysis of the sugar moiety leads to a complex picture. For example, epimerization at the 2′ position, *i*.*e*., converting Vidarabine to Adenosine, increases the compound’s activity at lower concentrations but leads to a loss of activity at higher concentrations. Substitution of the sugar 3′ – OH with a – H, found in Cordycepin, also hampers activity. Interestingly, for Clofarabine and Cladribine, the scenario is entirely different. Inclusion of a different halogen atom (*e*.*g*., Cl) in the 2′ position of the purine ring with concomitant fluorination (in Clofarabine) or reduction (in Cladribine) at the sugar 2′ completely abolishes the anti-viral activity. These observations illuminated the importance of purine ring – NH2 at R1 and sugar – OH at 2′ for the anti-viral activity of the drugs.

We concluded the structural analysis by examining the combination outcome from each of the nine Vidarabine derivatives with each of the eight Theobromine analogs. Strikingly, out of all 72 combinations, the only potent combination is found between Arainosine (or its prodrug Vidarabine) and Theobromine. Hence, while several nucleoside analogs were just as potent individually, if not more, than Vidarabine, none exhibited synergism. Similarly, none of the Theobromine analogs were active synergistically, even though they differed by only a single methyl group.

Clearly, detailed structural studies will be needed to explain the remarkable activity of Theobromine and Arainosine. However, based on the results presented in this study, it is possible to speculate as follows (Fig. 14): The activities of Theobromine and aminoadamantanes are similarly abolished by adamantanol (Fig. 8), pointing to a shared binding site. Moreover, Theobromine is significantly smaller than amantadine and, therefore, can still bind and inhibit the aminoadamantane-resistant S31N M2 channel, albeit at a substantially lower affinity than the aminoadamantane-sensitive channel. Moreover, against aminoadamantane-sensitive strains, Theobromine is individually less effective than aminoadamantanes, perhaps because its smaller size can only partially block proton transport. In contrast, adamantanol reduces the activity of Arainosine only moderately, but abolishes the activity of the duo entirely. Hence, it is possible to propose that Arainosine’s binding site only partially overlaps with the binding site for aminoadamantanes.

**Fig 14.**
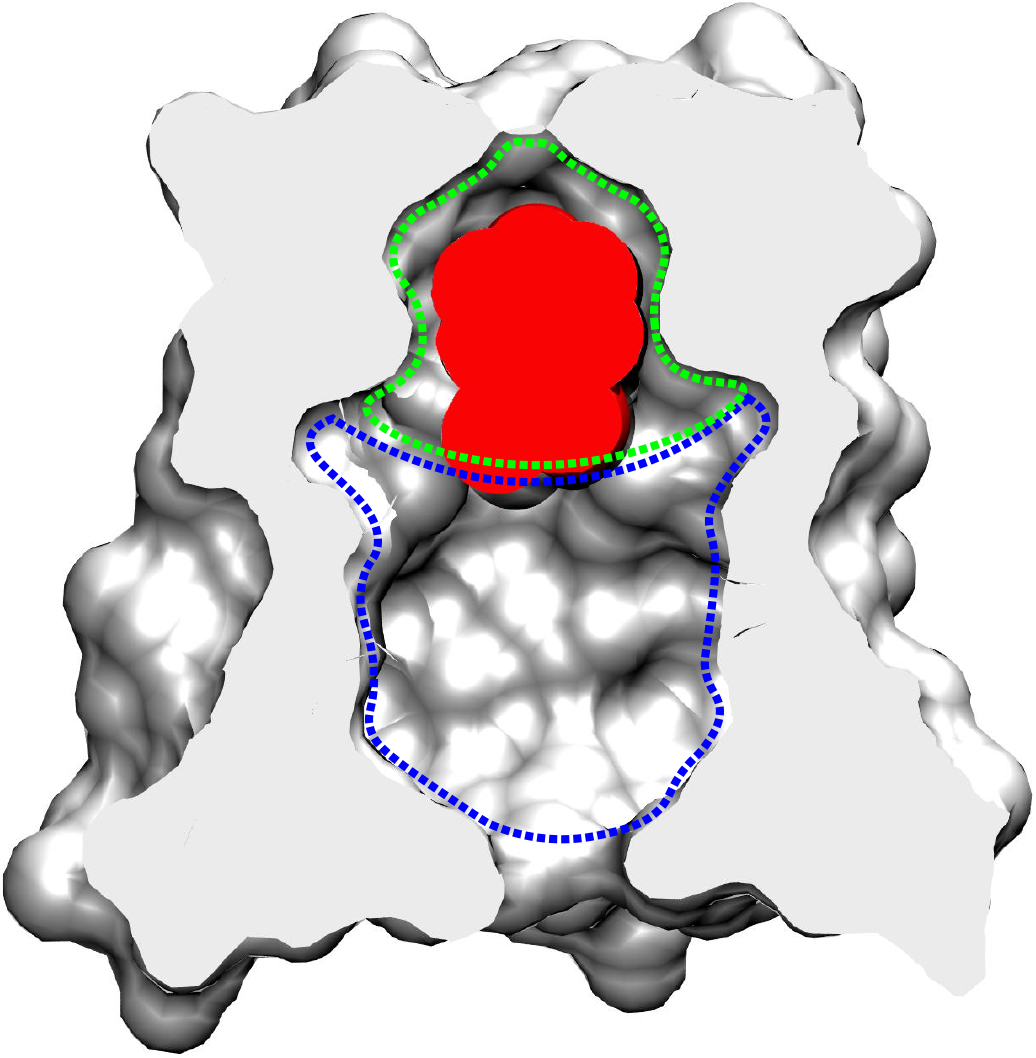
Proposed mechanism for inhibition of the Arainosine-Theobromine drug duo. The structure of the M2 protein bound to rimantadine (PDBID 6US9, (39)) in red is sliced in the middle (in gray) to depict the channel’s pore and drug binding site. Putative binding sites for Theobromine and Arainosine are shown in green and blue, respectively.

Having established the potent synergistic activity of Theobromine and Arainosine, we proceeded to examine the efficacy of the drug duo in an animal model. The *in-vivo* experiment on mice demonstrated the combined compounds’ activity in two viral H1N1 pandemic strains: pdm09 and mouse adaptive PR8. In the pdm09 strain, we did not observe significant disease symptoms such as weight loss in the case of the placebo. However, viral load quantification provided evidence of a clear reduction at two dosages of the drug duo: 10 mg/kg Theobromine with 15 mg/kg Arainosine and 3 mg/kg Theobromine with 4.5 mg/kg Arainosine (Fig. 11b). Oseltamivir at 20 mg/Kg exhibited a more minor reduction in viral load, which was not statistically significant. Hence, encouragingly, the results in tissue culture reproduced themselves in an animal model, whereby the drug duo of Theobromine and Arainosine outperforms Oseltamivir, even though it is given at a lower dosage.

Results employing the more virulent PR8 strain were even more encouraging. In this instance, dramatic differences in viral RNA load were seen in all treatments relative to the placebo (Fig. 11c). In particular, a dose-dependent reduction was seen in the drug duo, whereby with 10 mg/kg Theobromine and 15 mg/kg Arainosine, the reduction approached 16,000-fold relative to the placebo. Oseltamivir at 20 mg/Kg was also efficacious, yielding a ca. 3,000-fold reduction. Similar results were obtained when monitoring disease symptoms, whereby animals treated with the placebo required humane sacrifice four days post-infection. However, treatment with the drug duo was able to mitigate disease symptoms appreciably in a dose-dependent manner (Fig. 12). For example, oral treatment with 10 mg/kg of Theobromine with 15 mg/kg of Arainosine led to weight loss of only 6%. This result far exceeded the ability of Oseltamivir at 20 mg/Kg to reduce weight loss, which yielded an outcome of 15%.

## Conclusion

We present the results of a study aiming to identify new antiviral options against influenza. Our chosen route focused on a well-known drug target of the virus - its M2 H^+^ channel. Starting with a bacteria-based assay that screened a repurposed drug library, we identified a handful of potential blockers. These hits also served to search for similar compounds for further testing. Studies in tissue culture demonstrated that many of these compounds exhibit anti-viral activity. Moreover, combination studies yielded a remarkable synergy between Theobromine and Arainosine, which outperformed Oseltamivir, the leading drug against influenza. Interestingly, in an aminoadamantane-sensitive viral strain, the combination was efficacious, and the individual compounds were potent as well. Hence, resistivity against aminoadamantanes necessitated a combination of both drugs. Importantly, the drug duo exhibited anti-viral activity against two of the most important pandemic influenza strains: H1N1 swine flu and H5N1 avian flu. Finally, the tissue culture results were mirrored in an animal study that showed that the drug duo was particularly potent at lowering viral RNA and mitigating disease symptoms, far more so than Oseltamivir.

## ACKNOWLEDGMENTS

This work was supported in part by grants from the Israeli Science Foundation (948/19), The National Institutes of Health (1U19AI179421), and the Binational Agricultural Research and Development Fund (IS-5567-23 R). The following reagent was obtained through BEI Resources, NIAID, NIH: Influenza A Virus, A/Wisconsin/629-D02452/2009 (H1N1)pdm09, NR-19810. The authors thank Irina Skoda from the Department of Virology, Kimron Veterinary Institute, Ministry of agriculture, Israel, for the H5N1 A/Israel/975/2023 virus.

## Supporting figures and tables

**Fig S1.**
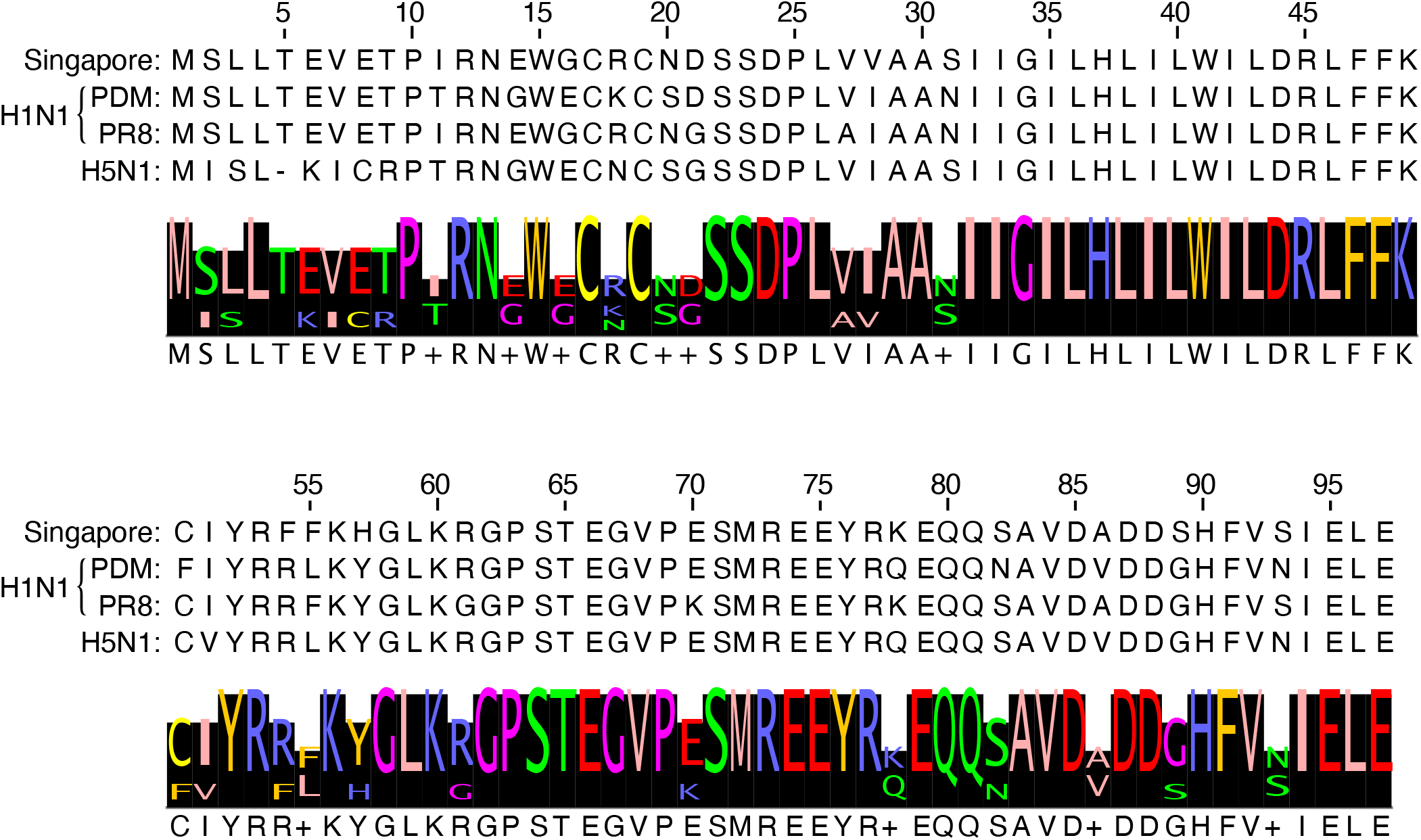
Sequences of the different M2 proteins used in the study. The sequence of the Singapore strain (A/Singapore/1/1957(H2N2)) was used in the bacteria-based screening, while the bottom three represent the M2 protein from the viruses used in the *in cellulo* and *in vivo* studies. The PDM strain is: A/Wisconsin/629-D02452/2009; the PR8 strain is: A/Puerto Rico/8/34; and the H5N1 strain is: A/Israel/975/2023

**Fig S2.**
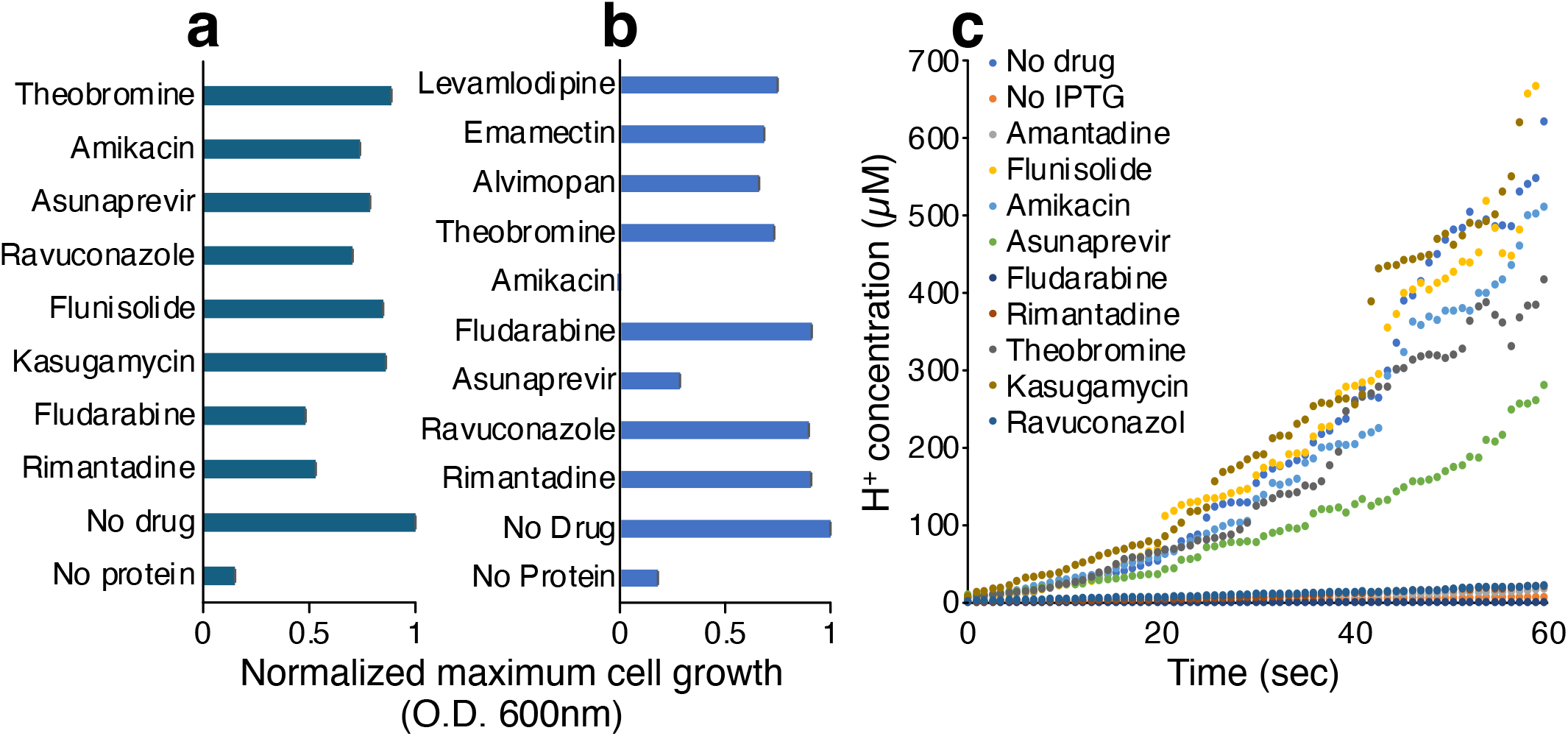
Bacteria-based positive assay to assess the activity of screened drugs against the aminoadamantane-sensitive M2 channel (a) and against a channel containing the S31N mutation (b). Panel c depicts the pH-dependent fluorescence assay on an aminoadamantane-sensitive M2 channel and the effects of drugs thereupon.

**Fig S3.**
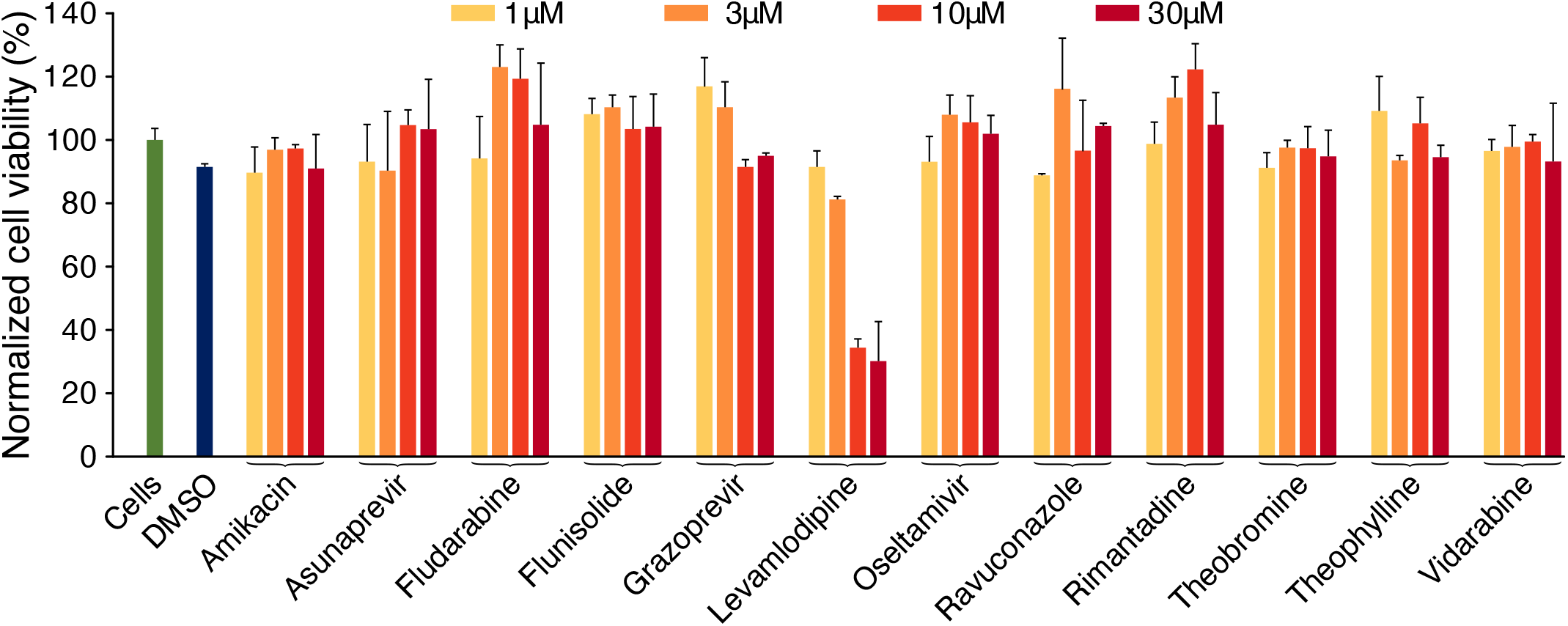
Toxicity of different compounds in tissue culture. Cellular viability after 48 hours was monitored by MTS as a function of different compounds at four concentrations, as noted. Untreated cells were used as a comparison and normalized to 100%. DMSO represents the vehicle.

**Fig S4.**
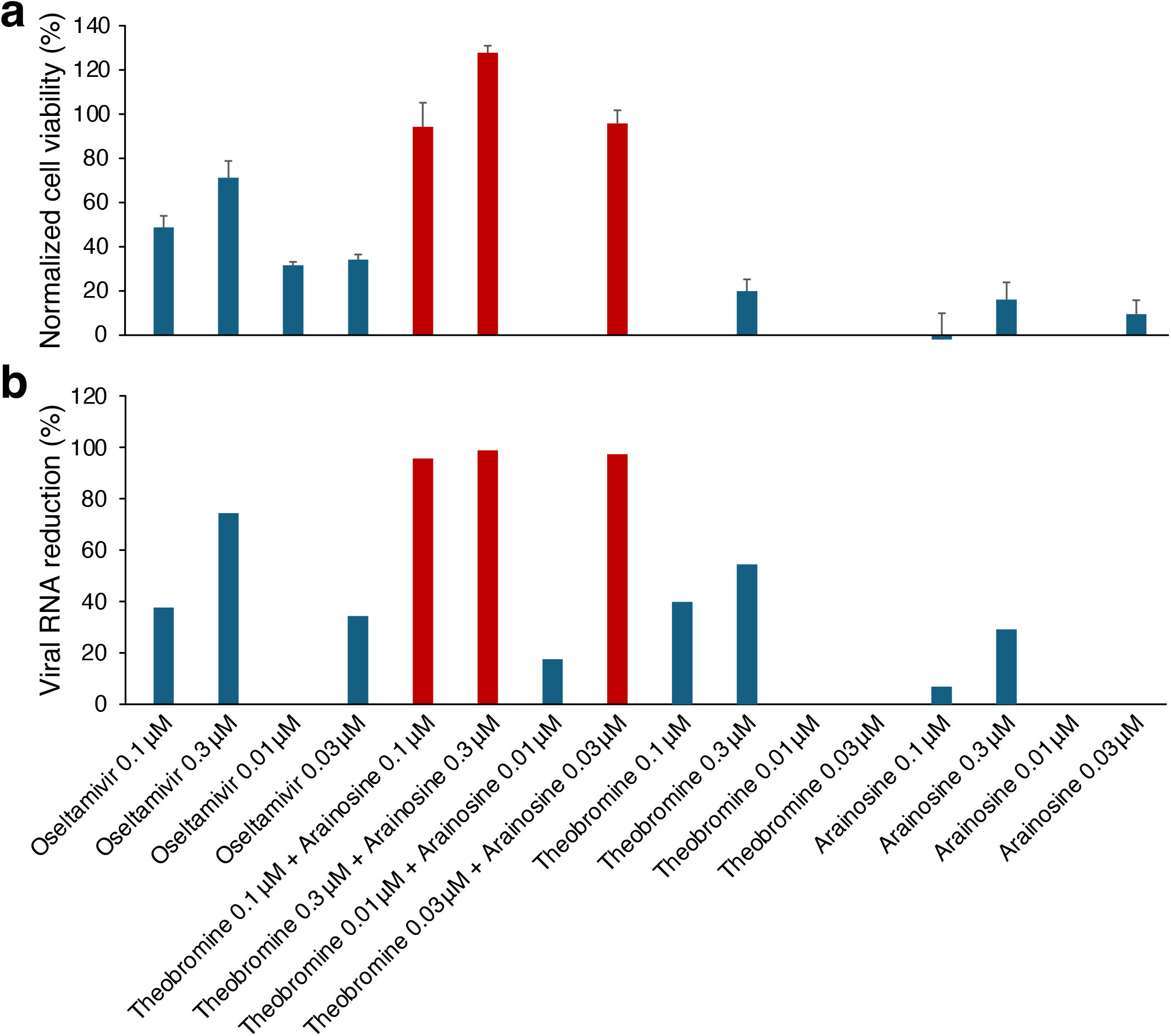
Antiviral activity (a) and viral load reduction (b) for different combinations of Theobromine and Arainosine on PR8 H1N1 Influenza strain.

**Fig S5.**
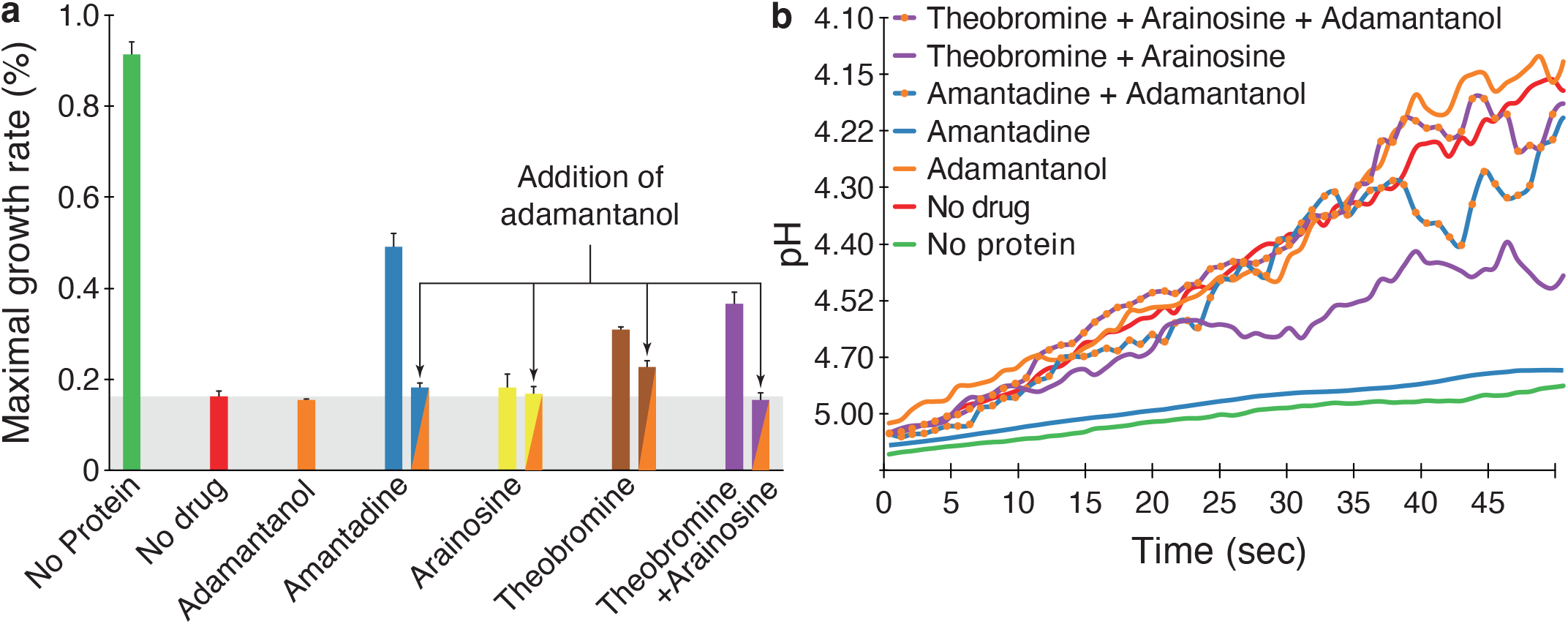
Effect of Adamantanol on the activities of Amantadine, Theobromine, Arainosine, and the Theobromine-Arainosine combination in the bacteria-based assays. (a) Negative assay in which bacterial growth is retarded by the M2 channel, and consequently blockers increase growth rate. (b) Fluorescence based assay, in which the M2 protein increases acidification, but effective blockers neutralize the channel’s activity. Note that Adamantanol reduces the activity of the drugs in both assays, except for Arainosine that exhibits a minor reduction.

**Table S1.** General mouse scoring table part I.

| Variable | Score | Description |
| --- | --- | --- |
| Appearance | 0 | Coat is smooth |
|  | 1 | Patches of hair piloerected |
|  | 2 | Majority of back is piloerected |
|  | 3 | Piloerection may or may not be present, hunched back |
|  | 4 | Piloerection may or may not be present, mouse appears emaciated |
| Level of consciousness | 0 | Mouse is active |
|  | 1 | Mouse is active but avoids standing upright |
|  | 2 | Mouse activity is noticeably slowed. The mouse is still ambulant. |
|  | 3 | Activity is impaired. Mouse only moves when provoked, movement have a tremor |
|  | 4 | Activity severely impaired. Mouse remains stationary when provoked, with possible tremor |
| Activity | 0 | Normal amount of activity. Mouse is any of: eating, drinking, climbing, running, fighting |
|  | 1 | Slightly suppressed activity. Mouse is moving around bottom of cage |
|  | 2 | Suppressed activity. Mouse is stationary with occasional investigative movements |
|  | 3 | No activity. Mouse is stationary |
|  | 4 | No activity. Mouse experiencing tremors, particularly in the hind legs |
| Response to stimulus | 0 | Mouse response immediately to auditory stimulus or touch |
|  | 1 | Slow or no response to auditory stimulus; strong response to touch (moves to escape) |
|  | 2 | No response to auditory stimulus; moderate response to touch (moves a few steps) |
|  | 3 | No response to auditory stimulus; mild response to touch (no locomotion) |
|  | 4 | No response to auditory stimulus; little or no response to touch. Cannot right itself if pushed over |

**Table S2.**
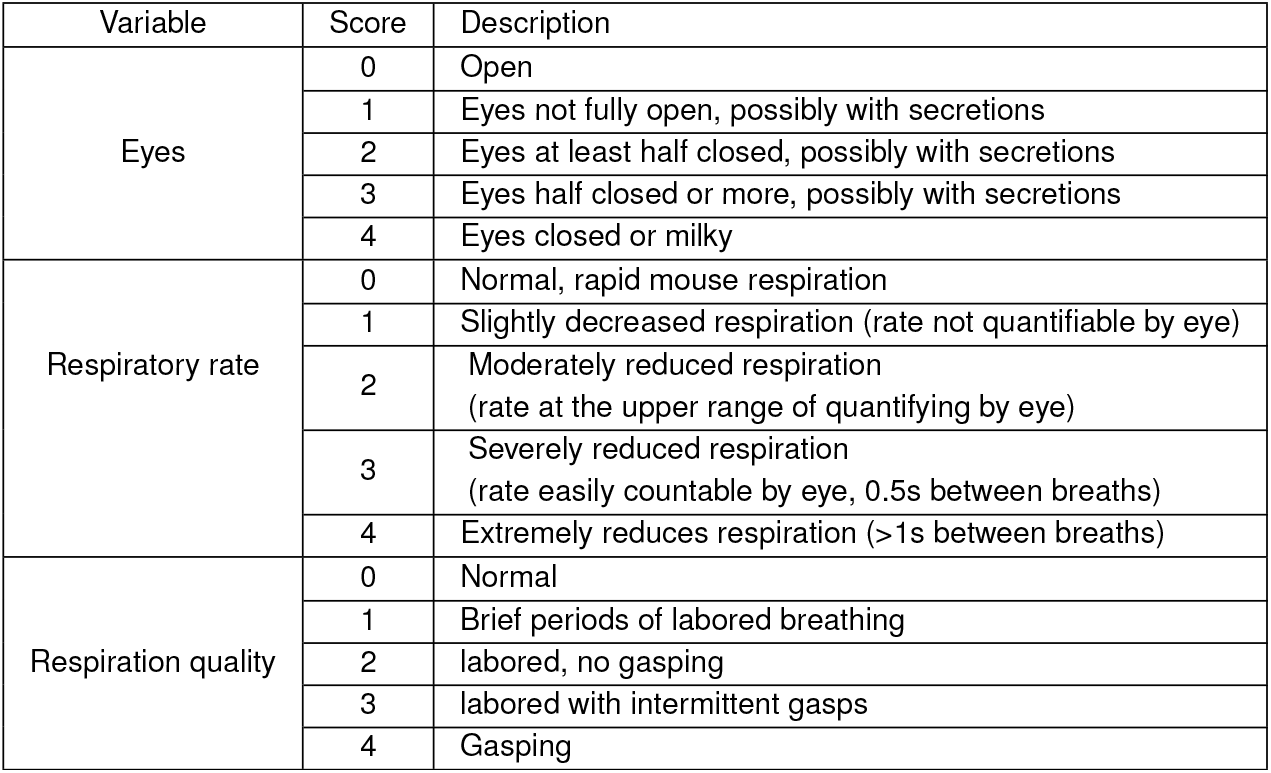
General mouse scoring table part II.

| Variable | Score | Description |
| --- | --- | --- |
| Eyes | 0 | Open |
|  | 1 | Eyes not fully open, possibly with secretions |
|  | 2 | Eyes at least half closed, possibly with secretions |
|  | 3 | Eyes half closed or more, possibly with secretions |
|  | 4 | Eyes closed or milky |
| Respiratory rate | 0 | Normal, rapid mouse respiration |
|  | 1 | Slightly decreased respiration (rate not quantifiable by eye) |
|  | 2 | Moderately reduced respiration (rate at the upper range of quantifying by eye) |
|  | 3 | Severely reduced respiration (rate easily countable by eye, 0.5s between breaths) |
|  | 4 | Extremely reduces respiration (>1s between breaths) |
| Respiration quality | 0 | Normal |
|  | 1 | Brief periods of labored breathing |
|  | 2 | labored, no gasping |
|  | 3 | labored with intermittent gasps |
|  | 4 | Gasping |

## Bibliography

1. M Maurel, et al., Interim 2023/24 influenza a vaccine effectiveness: Vebis european primary care and hospital multicentre studies, september 2023 to january 2024. Euro Surveill 29 (2024).

2. AD Iuliano, et al., Estimates of global seasonal influenza-associated respiratory mortality: a modelling study. Lancet 391, 1285–1300 (2018).

3. KD Kochanek, J Xu, E Arias, Mortality in the united states, 2019. NCHS Data Brief, 1–8 (2020).

4. RE Shope, The etiology of swine influenza. Science 73, 214–5 (1931).

5. RE Shope, Swine influenza : I. experimental transmission and pathology. J Exp Med 54, 349–59 (1931).

6. PA Lewis, RE Shope, Swine influenza : Ii. a hemophilic bacillus from the respiratory tract of infected swine. J Exp Med 54, 361–71 (1931).

7. RE Shope, Swine influenza : Iii. filtration experiments and etiology. J Exp Med 54, 373–85 (1931).

8. W Smith, C Andrewes, P Laidlaw, a virus obtained from influenza patients. The Lancet 222, 66–68 (1933) Originally published as Volume 2, Issue 5732.

9. S Tong, et al., A distinct lineage of influenza a virus from bats. Proc Natl Acad Sci U S A 109, 4269–74 (2012).

10. S Chakraborty, A Chauhan, Fighting the flu: a brief review on anti-influenza agents. Biotechnol Genet. Eng Rev 40, 858–909 (2024).

11. W Davies, et al., Antiviral activity of 1-adamantanamine (amantadine). Science 144, 862–863 (1964).

12. A Hay, N Kennedy, J Skehel, G Appleyard, The matrix protein gene determines amantadine-sensitivity of influenza viruses. J. Gen. Virol. 42, 189–191 (1979).

13. LH Pinto, LJ Holsinger, RA Lamb, Influenza virus m2 protein has ion channel activity. cell 69, 517–528 (1992).

14. A Kolocouris, et al., Aminoadamantanes with persistent in vitro efficacy against h1n1 (2009) influenza a. J. medicinal chemistry 57, 4629–4639 (2014).

15. ME Gonzalez, L Carrasco, Viroporins. FEBS Lett 552, 28–34 (2003).

16. D Assa, R Alhadeff, M Krugliak, IT Arkin, Mapping the resistance potential of influenza’s h+ channel against an antiviral blocker. J. molecular biology 428, 4209–4217 (2016).

17. A Hay, A Wolstenholme, J Skehel, MH Smith, The molecular basis of the specific anti-influenza action of amantadine. The EMBO journal 4, 3021–3024 (1985).

18. P Astrahan, I Kass, MA Cooper, IT Arkin, A novel method of resistance for influenza against a channel-blocking antiviral drug. Proteins 55, 251–7 (2004).

19. S Stumpe, EP Bakker, Requirement of a large K<sup>+</sup>-uptake capacity and of extracytoplasmic protease activity for protamine resistance of escherichia coli. Arch. microbiology 167, 126–136 (1997).

20. P Santner, et al., A robust proton flux (phlux) assay for studying the function and inhibition of the influenza a m2 proton channel. Biochemistry 57, 5949–5956 (2018).

21. G Miesenböck, DA De Angelis, JE Rothman, Visualizing secretion and synaptic transmission with ph-sensitive green fluorescent proteins. Nature 394, 192–195 (1998).

22. T McIlvaine, A buffer solution for colorimetric comparison. J. biol. Chem 49, 183–186 (1921).

23. P Astrahan, et al., Quantitative analysis of influenza m2 channel blockers. Biochimica et Biophys. Acta (BBA)-Biomembranes 1808, 394–398 (2011).

24. M Hussain, HD Galvin, TY Haw, AN Nutsford, M Husain, Drug resistance in influenza a virus: the epidemiology and management. Infect Drug Resist. 10, 121–134 (2017).

25. W Shen, et al., 5′-od-valyl ara. a, a potential prodrug for improving oral bioavailability of the antiviral agent vidarabine. Nucleosides, Nucleotides Nucleic Acids 28, 43–55 (2009).

26. H Leonov, P Astrahan, M Krugliak, IT Arkin, How do aminoadamantanes block the influenza m2 channel, and how does resistance develop? J Am Chem Soc 133, 9903–11 (2011).

27. WW Lee, A Benitez, L Goodman, BR Baker, Potential anticancer agents.1 xl. synthesis of the β-anomer of 9-(d-arabinofuranosyl)-adenine. J. Am. Chem. Soc. 82, 2648–2649 (1960).

28. T Francis Jr, Transmission of influenza by a filterable virus. Science 80, 457–459 (1934).

29. K Basu, M Krugliak, IT Arkin, Viroporins of mpox virus. Int. J. Mol. Sci. 24, 13828 (2023).

30. H Lahiri, IT Arkin, Searching for blockers of dengue and west nile virus viroporins. Viruses 14, 1750 (2022).

31. R Taube, R Alhadeff, D Assa, M Krugliak, IT Arkin, Bacteria-based analysis of hiv-1 vpu channel activity. PLoS One 9, e105387 (2014).

32. PPS Tomar, R Oren, M Krugliak, IT Arkin, Potential viroporin candidates from pathogenic viruses using bacteria-based bioassays. Viruses 11, 632 (2019).

33. PPS Tomar, M Krugliak, IT Arkin, Blockers of the sars-cov-2 3a channel identified by targeted drug repurposing. Viruses 13, 532 (2021).

34. PPS Tomar, M Krugliak, A Singh, IT Arkin, Zika m—a potential viroporin: Mutational study and drug repurposing. Biomedicines 10, 641 (2022).

35. CU Kim, et al., Influenza neuraminidase inhibitors possessing a novel hydrophobic interaction in the enzyme active site: design, synthesis, and structural analysis of carbocyclic sialic acid analogues with potent anti-influenza activity. J Am Chem Soc 119, 681–90 (1997).

36. P Jaaks, et al., Effective drug combinations in breast, colon and pancreatic cancer cells. Nature 603, 166–173 (2022).

37. I Shumilin, C Allolio, D Harries, How sugars modify caffeine self-association and solubility: Resolving a mechanism of selective hydrotropy. J Am Chem Soc 141, 18056–18063 (2019).

38. F Fauvelle, et al., Pharmacokinetics of vidarabine in the treatment of polyarteritis nodosa. Fundam Clin Pharmacol 6, 11–5 (1992).

39. JL Thomaston, et al., Rimantadine binds to and inhibits the influenza a m2 proton channel without enantiomeric specificity. Biochemistry (2021).

